# AI-aided chronic mixture risk assessment along a small European river reveals multiple sites at risk and pharmaceuticals being the main risk drivers

**DOI:** 10.1101/2024.11.15.623722

**Authors:** Fabian G. Weichert, Pedro A. Inostroza, Jörg Ahlheim, Thomas Backhaus, Werner Brack, Mario Brauns, Patrick Fink, Martin Krauss, Patrik Svedberg, Henner Hollert

## Abstract

The vast amount of registered chemicals leads to a high diversity of substances occurring in the environment and the creation of new substances outpaces chemical risk assessment as well as monitoring strategies. Hence, risk assessment strategies need to be modified ensuring that they remain aligned with the rapid development and marketing of new substances. Here we performed a longitudinal chronic mixture risk assessment considering a real-world case study scenario with diverse anthropogenic impact types characterised by different land uses along a river in Central Germany. We sampled river water using large-volume solid phase extraction at six selected sampling sites. Following chemical analysis using liquid chromatography-high resolution mass spectrometry, we quantified 192 substances. For 34% of them, we obtained empirical chronic effect data for freshwater organisms. Furthermore, we used the open-source artificial intelligence (AI) model TRIDENT to predict chronic toxicity for all substances. A multi-scenario mixture risk assessment was conducted for three taxonomic groups, using the concentration-addition concept and considering various hazard and exposure scenarios. The results showed that the chronic risk estimates for all taxonomic groups were considerably higher when the empirical data was amended with data from in silico modelling. We identified hot spots of chemical pollution and our analysis indicated that fish were the most vulnerable taxonomic group, with pharmaceuticals being the most relevant risk drivers. Our study exemplifies the application of an AI model to predict chronic risk for aquatic organisms in combination with the consideration of multiple risk scenarios, that may complement future risk assessment strategies.

**Highlights:**

- 192 organic chemicals were quantified in six surface water samples along a river.
- Multiple hazard and exposure scenarios were considered in mixture risk assessment.
- Artificial intelligence was used to fill data gaps and predict chronic ecotoxicity.
- Fish were identified as the most vulnerable taxonomic group for chronic toxicity.
- Pharmaceuticals were the most prevalent mixture risk drivers.

**Graphical abstract:** 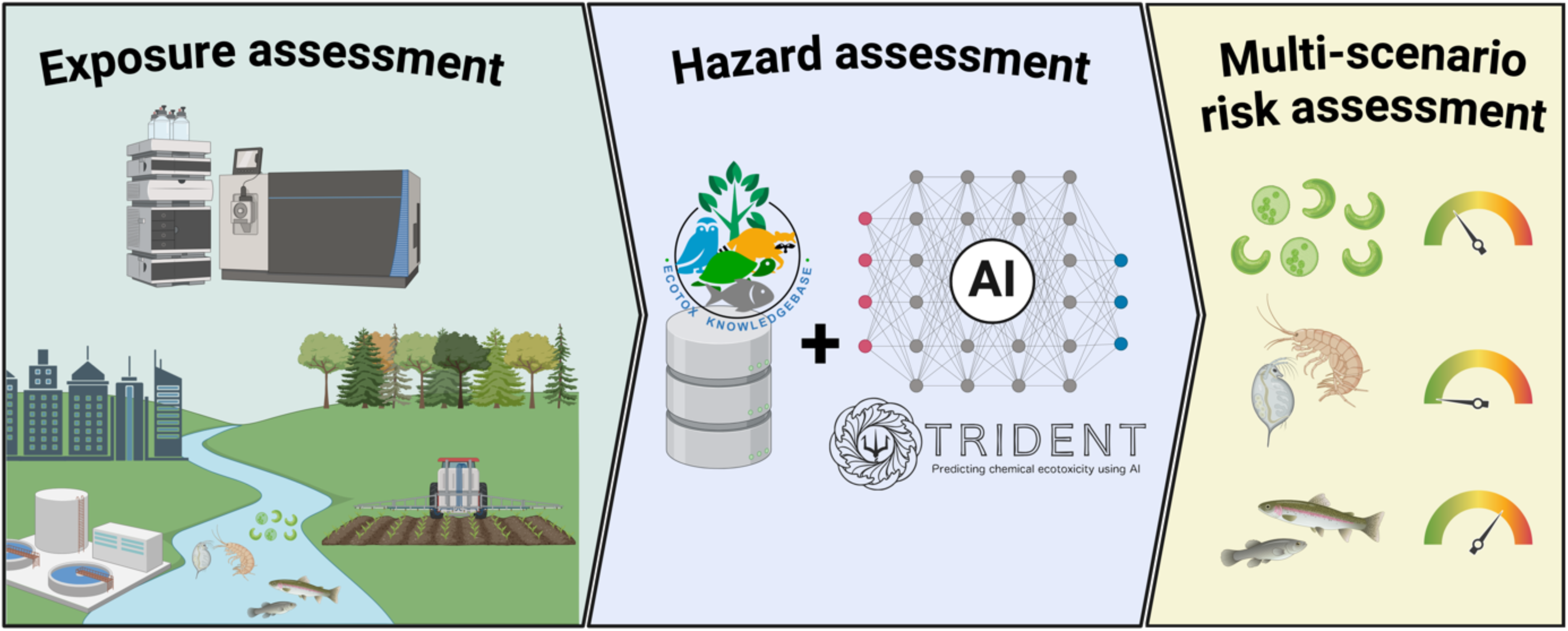

## 1. Introduction

Environmental pollution is one of the five main drivers of global biodiversity loss (IPBES, 2019) and a major driver of global change (Bernhardt et al., 2017). Furthermore, it is a major contributor to the deteriorated ecological status of surface waters (Lemm et al., 2021; Malaj et al., 2014). However, the consequences for biodiversity and functions of ecosystems are not fully understood (Sylvester et al., 2023; Wang et al., 2020). There are approximately 350,000 different chemical substances and mixtures currently registered for use and production (Wang et al., 2020). In addition, more than 30 new chemicals are discovered every hour (Backhaus et al., 2012), some of which will likely be incorporated into new industrial and consumer products in the future. This rate of chemical discovery far outpaces the capacity for comprehensive chemical risk assessment and monitoring (Persson et al., 2022; Richardson et al., 2023; Wang et al., 2020). Consequently, new risk assessment tools and strategies are required with the capabilities and capacities to keep up with the rapid development and marketing of new substances.

Traditionally, risk assessment has focused on individual substances, largely ignoring mixture effects. Within the last decade, awareness of mixture risks has been growing (Drakvik et al., 2020; Treu et al., 2024), which is also reflected in the European Union’s Chemicals Strategy for Sustainability (European Commission, 2020).

Mixture risk assessment (MRA) strategies are often based on the concentration addition (CA) assumption, which has been shown to be a reliable estimate and well applicable to environmental mixtures with known components (Backhaus and Faust, 2012; Gustavsson et al., 2017; Inostroza et al., 2024). However, CA requires information on the (eco)toxicity of all mixture components, which is a major hurdle for its application to complex environmental mixtures (Finckh et al., 2022; Gustavsson et al., 2023; Spilsbury et al., 2020). Achieving a comprehensive risk assessment of real-world exposure scenarios is therefore not feasible without the integration of *in silico* techniques for predicting toxicities, such as Quantitative Structure-Activity Relationships (QSARs), in order to cope with the sheer number of substances lacking empirical (eco)toxicity data.

Classical QSAR models largely build on regression models that correlate structural features and physicochemical properties, such as the log *K*_OW_ of a substance, with its activity (toxicity). Even though QSAR models have been used in ecotoxicological research for decades, their use has been critically discussed. For instance, challenges include diverse and heterogeneous datasets introducing noise and inconsistencies into models. Also, there are unreliable predictions for compounds that fall outside the narrow applicability domains of several classical QSAR models (De et al., 2022; Gramatica, 2013).

A new generation of (eco)toxicity modelling techniques has therefore begun to gain traction. Artificial Intelligence (AI) and deep learning methodologies have been successfully used to predict e.g., molecular functioning (Zhang et al., 2023) as well as acute toxicity (Wu et al., 2022; Zhang et al., 2022; Zubrod et al., 2023), and the scientific literature on the potential uses of AI is growing (Kleinstreuer and Hartung, 2024; Rillig et al., 2024; von Borries et al., 2023). Very recent advances in the field show that transformers in connection with deep neural networks have been shown to deliver remarkably accurate predictions of both acute and chronic toxicity in aquatic organisms (Gustavsson et al., 2024). These new tools have great potential to improve the risk assessment of complex environmental mixtures and so far, no deep learning-based approaches to toxicity prediction have been used in chronic MRA to the best of our knowledge.

The present study therefore implements a comprehensive AI-aided longitudinal chronic risk assessment in a real-world scenario, considering various anthropogenic impact types along different river stretches. In this study, we identify (1) hot spots of chemical pollution to isolate priority sites for further investigation, (2) the taxonomic groups that are particularly vulnerable to the mixtures detected at the sampling sites, and (3) the risk drivers amongst the multitude of encountered chemicals. A major focus of this paper is the use of different exposure and hazard scenarios to account for uncertainty due to chemicals present below chemical-analytical detection limits and the uncertainty of the predicted ecotoxicity data.

## 2. Materials & methods

Detailed descriptions of the case study area, the sampling sites, sampling procedures, sample preparation as well as liquid chromatography-high resolution mass spectrometry (LC-HRMS) analysis of organic micropollutants are published in Weichert et al. (2024a). Therefore, only brief summaries are given below.

### 2.1. Case study area

The Holtemme River (length: ca. 47 km; catchment area: ca. 282 km^2^) in Central Germany is part of the Elbe/Saale/Bode river system and exemplifies a low mountain stream with a gradient of anthropogenic influence (Fink et al., 2020). Sources of chemical pollution include run-off from agricultural land or urban areas, rainwater retention basins as well as two wastewater treatment plants, Silstedt and Halberstadt, with 80,000 and 60,000 population equivalents, respectively (Bartels et al., 2020; Landesamt für Umweltschutz Sachsen-Anhalt, 2019; Weitere et al., 2021). Within the last decade, the Holtemme River has been a site of interest for various studies, including chemical exposure analysis in different matrices, biological effect assessment of various *in vitro* and *in vivo* endpoints and their interconnection (Beckers et al., 2020, 2018; Inostroza et al., 2017, 2016; Muschket et al., 2021, 2018; Schmitz et al., 2022; Švara et al., 2021; Weichert et al., 2024a; Weitere et al., 2021), which is of great value for the prioritisation of the sampling sites used in the present study.

### 2.2. Sites, sampling and sample preparation

In October 2021, six sampling sites along the river course (**Site1, Site2, Site3, Site4, Site5** and **Site6**) were selected, based on knowledge from prior studies (Beckers et al., 2020, 2018; Inostroza et al., 2017, 2016; Schmitz et al., 2022; Švara et al., 2021; Weitere et al., 2021), covering distinct land uses (Figure 1, Table S1) and therefore various anthropogenic impact types. At each sampling site, 25 L of river water were sampled by large-volume solid phase extraction (LVSPE) (Schulze et al., 2017). In addition to the samples collected at the above sampling locations, a trip blank sample cartridge was carried under the same conditions during the field trip and eluted in the same manner as the samples to provide a control for potential processing-related contamination (Weichert et al., 2024a).

**Figure 1.**
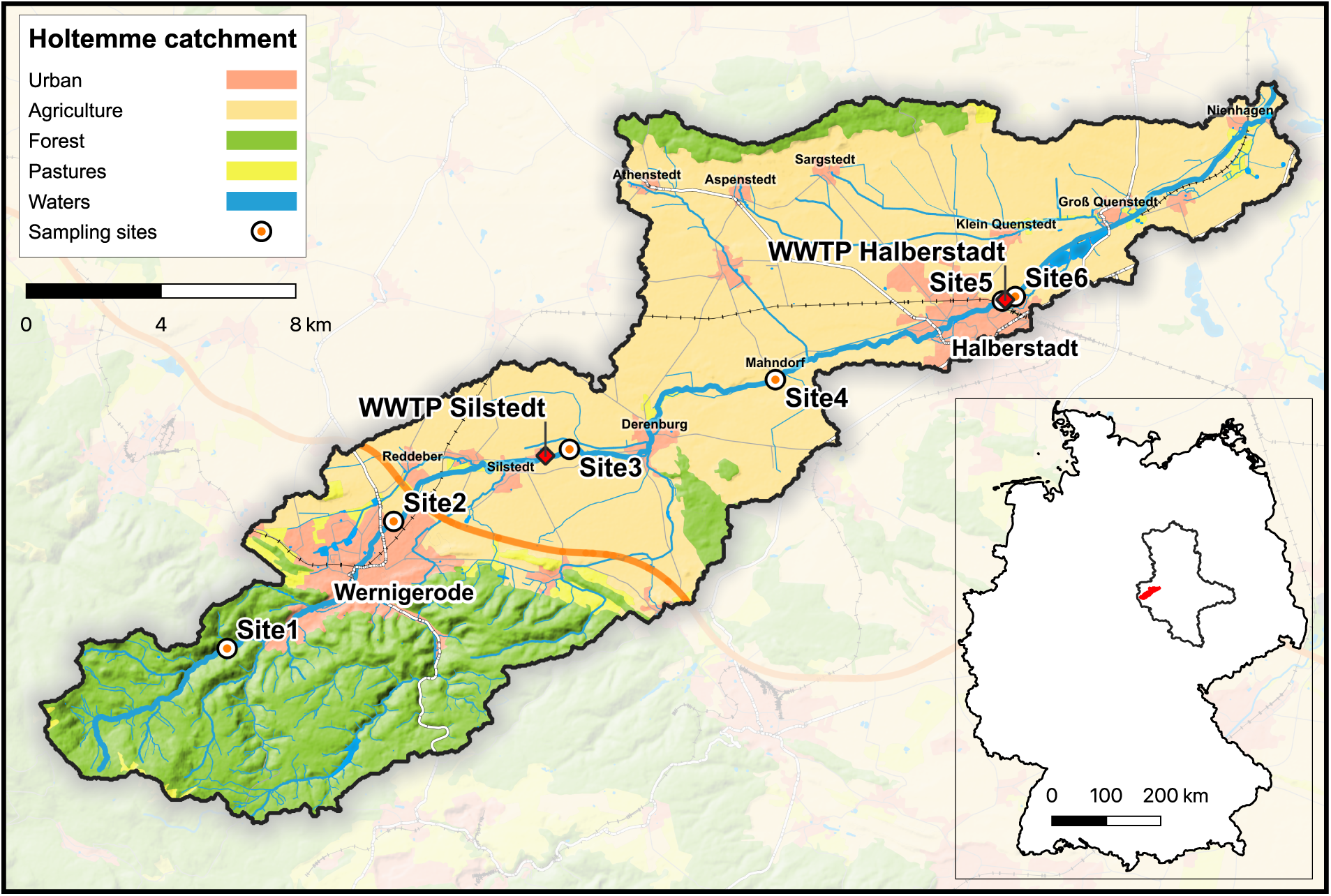
Representation of the Holtemme River catchment, showing the sampling sites, wastewater treatment plants and land-use within the area. Generated using European Union’s Copernicus Land Monitoring Service information; https://doi.org/10.2909/71c95a07-e296-44fc-b22b-415f42acfdf0.

### 2.3. LC-HRMS analysis

Briefly, reconstituted LVSPE extracts (with a relative enrichment factor (REF) of 1000) were spiked with an internal standard mixture consisting of 39 isotope-labelled compounds (Finckh et al., 2022) and injected into an UltiMate 3000 LC system (Thermo Scientific). After separation on a reversed-phase column, the samples were analysed with a QExactive Plus quadrupole-Orbitrap HRMS (Thermo Scientific) using electrospray ionisation (ESI) in positive and negative mode. Finally, the R package {MZquant} (Schulze et al., 2021) was used to quantify the target compounds after blank correction and calibration. The method detection limits (MDLs) for each target compound were determined based on the method detailed in (US EPA, 2016).

### 2.4. Empirical ecotoxicological data

Empirical ecotoxicological data were retrieved based on the descriptions in Inostroza et al. (2024). Briefly, data were obtained from the US EPA ECOTOXicology Knowledgebase (ECOTOX, version ecotox_ascii_09_14_2023, (Olker et al., 2022). Only freshwater data were retained, and records lacking essential information such as exposure durations, measurement endpoints, concentration units, or valid data sources, as well as those that only report limit values, were excluded. Following the recommendations of Warne et al. (2018), only chronic data was used (exposure durations of > 1, ≥ 14, and ≥ 21 days for algae, aquatic invertebrates, and fish, respectively) and recalculated to chronic EC_10_-equivalents (NOEL/NOECs and EC/IC/LC_1-20_-values, MATCs, LOEL/LOECs, EC/ED/IC/LC_50_-values^1^ were divided by 1, 2, 2.5 and 5, respectively). The EC_10_-equivalents were converted to micromoles per litre (μmol/L), and taxonomic groups were assigned to species using an in-house dataset derived from reported phyla in ECOTOX. Finally, the geometric mean for each taxonomic group and chemical was calculated, to account for variation and provide a robust measure less susceptible to outliers compared to arithmetic means.

### 2.5. In silico predictions of chronic aquatic ecotoxicity

The Ecological Structure Activity Relationships (ECOSAR) Program (version 2.2, US EPA, Washington, DC, USA) was used to predict the chronic toxicity for green algae, daphnids (as representative of aquatic invertebrates), and fish for all chemicals detected. Chemical identification was based on corresponding canonical Simplified Molecular-Input Line-Entry System (SMILES) codes, which served as input to predict toxicity in the form of chronic values (ChV). When multiple predictions per taxonomic group and chemical were provided by the software, the geometric mean of the ChVs from different QSAR models was calculated. Lastly, the predicted ChVs were converted to μmol/L. More information on the methodologies used to obtain and consolidate ECOSAR data is detailed in Svedberg et al. (2023).

The TRIDENT model used to predict chronic toxicity is a *Bidirectional Encoder Representations from Transformers* (BERT)-based language model (ChemBERTa) combined with a deep neural network. The model and its performance are described in detail elsewhere (Gustavsson et al., 2024). Briefly, the TRIDENT transformer is pre-trained on a large dataset, comprising approximately 144k empirical toxicity measurements for 6469 substances in 1842 species. The model converts SMILES codes into a high-dimensional embedding vector which characterises the toxicity of the molecular structure. This vector is complemented with details on exposure duration (24 – 720 h), the toxicological effect of interest (mortality, growth, intoxication, reproduction or population effects) and endpoint (EC_10_ or EC_50_) before it is fed into a deep neural network. The output of this network is a predicted effect concentration (EC_50_ or EC_10_), for algae, aquatic invertebrates or fish.

All source codes and background information are located in online repositories (https://github.com/StyrbjornKall/TRIDENT and https://huggingface.co/StyrbjornKall) and the model can be used through a web interface (https://trident.serve.scilifelab.se/). The datasets for this study were obtained from the TRIDENT web interface. The batch of canonical SMILES codes for the 457 substances reliably measured during the study (Weichert et al., 2024b, 2024a) were uploaded and EC_10_-values were obtained for each taxonomic group. For algae, a dataset for the available toxicological effect *population* was retrieved (exposure duration: 24 h). For aquatic invertebrates, one dataset per toxicological effect (*mortality*, *intoxication*, *reproduction* and *population effects*) was downloaded with an exposure duration setting of 336 h (14 days). Lastly, EC_10_-values for fish were obtained for the available toxicological effects of *mortality* and *growth* at a duration setting of 504 h (21 days). For aquatic invertebrates and fish, the geometric mean was calculated from the predictions for each chemical.

### 2.6. Mixture risk assessment

Briefly, the MRA was based on the CA concept and performed separately for the three taxonomic groups algae, aquatic invertebrates and fish as well as for the most sensitive taxonomic group (MST approach). The latter considers only the smallest EC_10_-values across the three aforementioned taxonomic groups per chemical and is, therefore, the most protective environmental risk assessment approach (Backhaus and Faust, 2012; Gustavsson et al., 2017).

To express the risk for a specific taxonomic group, the risk quotient (RQ_ΣTU_) was defined as follows:

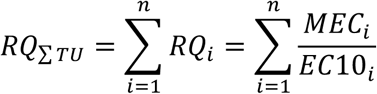

where *RQ_i_* is the risk quotient of chemical *i*, which is defined as the ratio between *MEC_i_* (measured environmental concentration) and *EC10_i_* (chronic EC_10_-equivalent) of chemical *i* for algae, aquatic invertebrates or fish. This reflects a summation of Toxic Units (TUs), thus dimensionless measures of the contribution of chemical *i* to the overall mixture toxicity.

For the MST approach, the mixture risk quotient (*RQ_MST_*) was defined as:

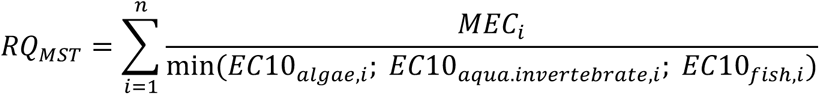

where *MEC_i_* is the measured environmental concentration of a chemical *i* and min(*EC10_algae,i_*; *EC10_aqua.invertebrate,i_*; *EC10_fish,i_*) is the smallest chronic EC_10_-equivalent of a chemical *i* for algae, aquatic invertebrates or fish.

For the calculation of RQs, three different exposure scenarios (ES) were defined based on the treatment of non-detects in MECs (Table 1).

**Table 1.** Description of handling of non-detects in the data set for individual exposure scenarios.

| Exposure scenarios | Handling of non-detects for the respective scenarios for the mixture risk assessment |
| --- | --- |
| ES1 | Non-detects are set to zero, representing the lowest risk scenario compatible with the data. |
| ES2 | Non-detects are set to their method detection limits (MDLs), representing the highest risk scenario compatible with the data. |
| ES3 | Non-detects are replaced by Kaplan-Meier estimate (Gustavsson et al., 2017; Helsel, 2005; Inostroza et al., 2024), providing an accurate basis for risk assessment but not allowing the identification of individual risk drivers. |

As summarised in Table 2, three main hazard scenarios (HS1, HS2, HS3) were used based on the toxicological data used. Two additional hazard scenarios were defined, based on only using the toxicological data from modelling approaches in ECOSAR and TRIDENT (HS4, HS5). We also included four specific hazard scenarios, assuming the modelling estimates may be off by two orders of magnitude (HS2a, HS2b, HS3a, HS3b). The idea of this approach is to consider the modelling uncertainty and is justified in more detail in Inostroza et al. (2024).

**Table 2.**
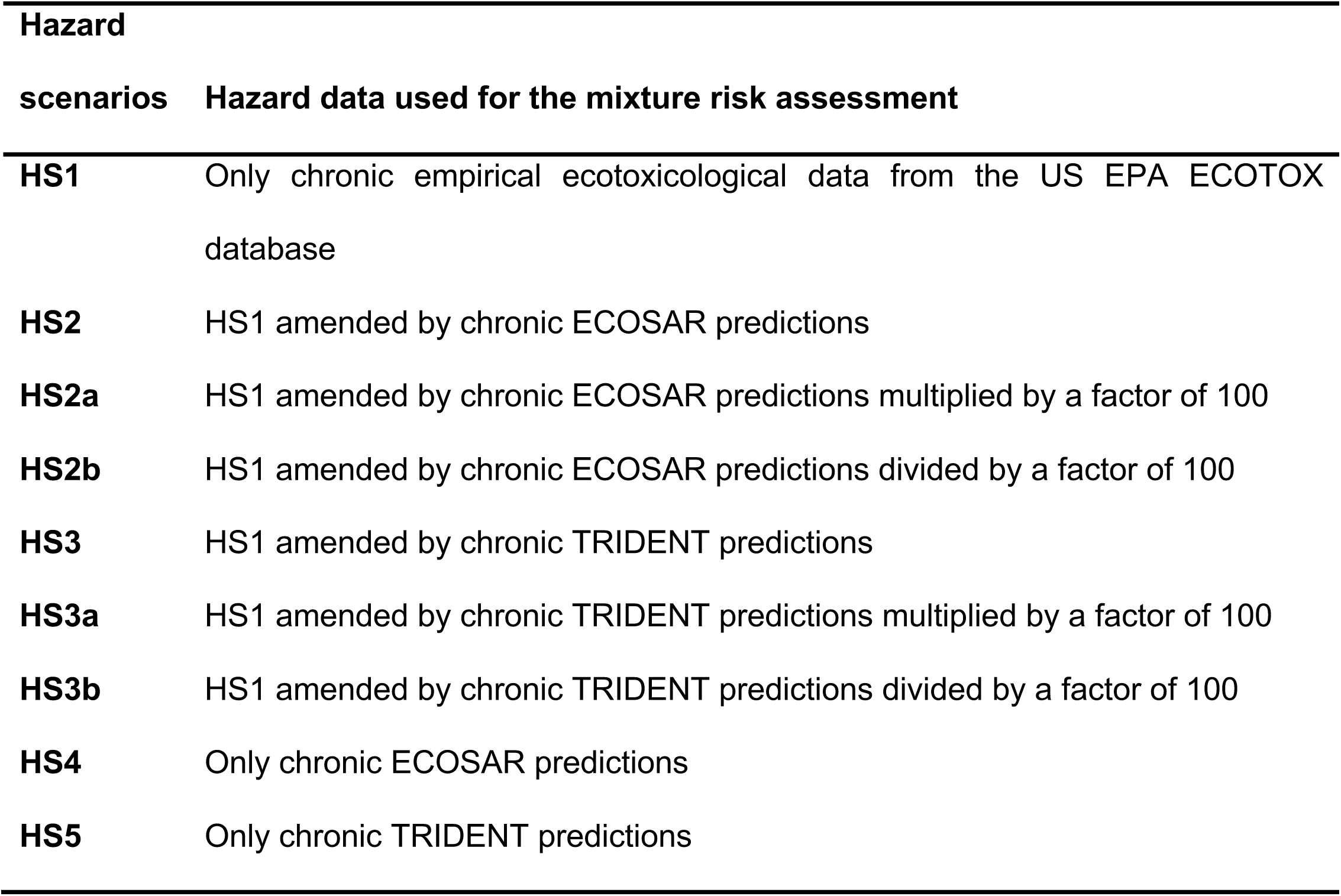
Description of the hazard data used for individual hazard scenarios.

From the ES1 and ES2, various types of mixture risk drivers were determined, as defined in Inostroza et al. (2024). Briefly, for the HS1 – HS3, ***actual*** mixture risk drivers were identified. These types of risk-driving substances are defined by their contribution to overall toxicity with an RQ_i_ ≥ 0.02. Additionally, ***potential*** mixture risk drivers were identified from the HS2b and HS3b. A substance is highlighted as a potential risk driver if the uncertainty of modelling approaches (two orders of magnitude below the predicted toxicity value) is considered and results in an RQ_i_ ≥ 0.02.

### 2.7. Data accessibility and data analysis

Data analyses and data visualisation were executed in R version 4.4.2 (R Development Core Team, 2008). The Kaplan-Meier estimators were calculated and integrated into the analysis using resources from the R-package {NADA} (Lee, 2020). Following the FAIR (Findable, Accessible, Interoperable, and Reusable) principles, all raw data including LVSPE measurements (Weichert et al., 2024b, 2024a), ECOTOX data, ECOSAR and TRIDENT predictions, and analysis scripts from R (Weichert et al., 2024c) are openly accessible through the platform Zenodo, ensuring transparency and reproducibility.

## 3. Results

### 3.1. Occurrence of organic chemicals along the Holtemme River

The chemical analysis data for the six sampling sites and the trip blank used in this study are published in a separate data article (Weichert et al., 2024a) and can be downloaded from the repository Zenodo (Weichert et al., 2024b). 192 substances from ten different chemical classes, from a total of 452 substances included in the analytical suite, were detected above the respective method detection limit (MDL) in at least one of the samples. In the trip blank sample and the samples from Site1 and Site2, only five, eight and 40 compounds were detected, respectively. In the samples from Site3, Site4, Site5 and Site6 we detected 157, 138, 126 and 169 individual compounds above the MDL (Table S2). A principal component analysis (PCA) of the concentrations found revealed that the samples from Site1 and Site2, situated in the upper reaches and within the city of Wernigerode, along with the trip blank, showed a comparable chemical fingerprint. Furthermore, samples from Site4 and Site5, collected between the two WWTPs, clustered closely, whereas samples from Site3 and Site6, located directly downstream of the WWTPs occupy a clearly different position in the PCA plot (Figure 2A).

**Figure 2.**
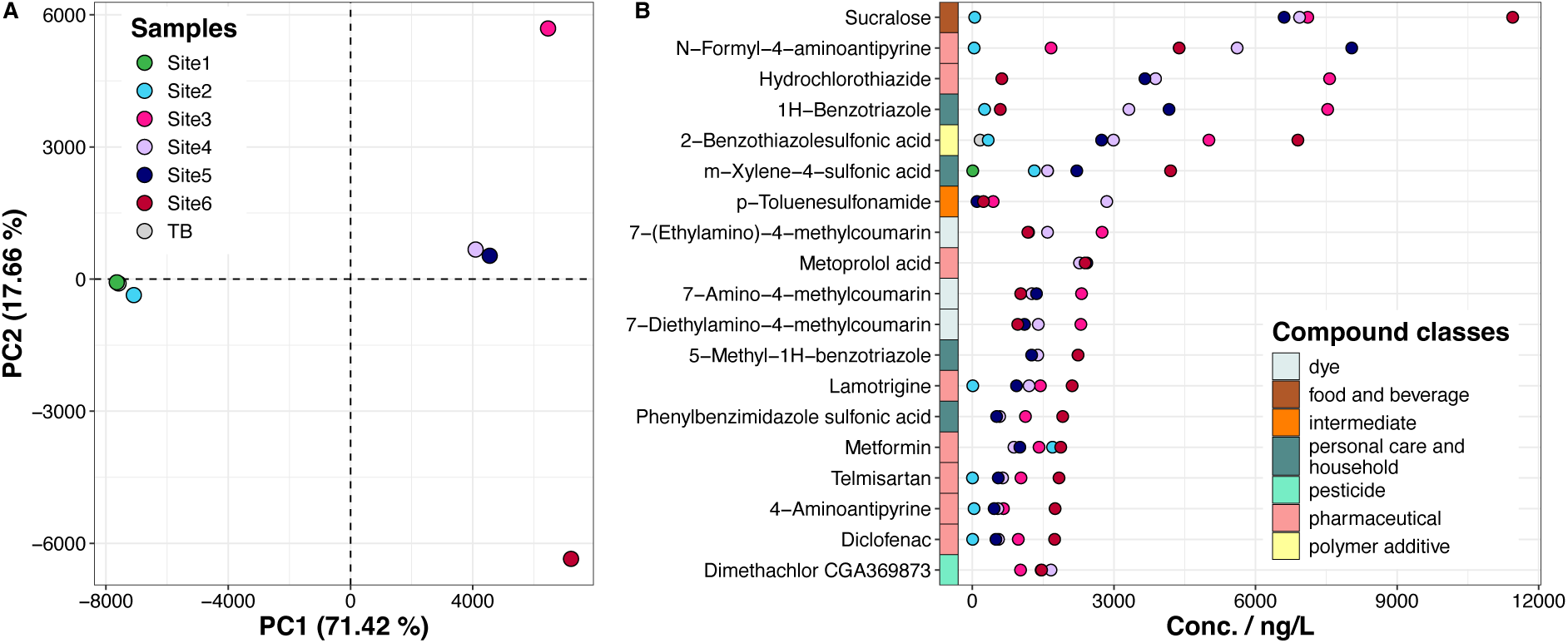
(A) Principal component analysis of the chemical profile for each sample (B) For each sample, the concentrations of the top 10% of compounds detected in at least one of the samples. The colours close to the compound names indicate the compound class as shown in panel B, while the colours of the dots represent the sampling site as indicated in panel A. TB: trip blank.

Considering the highest concentrations of each of the detected compounds, 19 substances from diverse chemical classes (from highest: sucralose, N-formyl-4-aminoantipyrine, hydrochlorothiazide, 1H-benzotriazole, 2-benzothiazolesulfonic acid, m-xylene-4-sulfonic acid, p-toluenesulfonamide, 7-(ethylamino)-4-methylcoumarin, metoprolol acid, 7-amino-4-methylcoumarin, 7-diethylamino-4-methylcoumarin, 5-methyl-1H-benzotriazole, lamotrigine, phenylbenzimidazole sulfonic acid, metformin, telmisartan, 4-aminoantipyrine, diclofenac, dimethachlor CGA369873) make up the top 10%, with concentrations ranging between 1.7 ng/L and 11.4 µg/L (Figure 2B).

### 3.2. Chronic effect data from the ECOTOX database and the *in silico* modelling

We retrieved empirical data for chronic toxicity from the ECOTOX database for 190 out of the total 457 target compounds. From the subset of the 192 compounds detected above the respective MDL at one or more sites, we retrieved data for 63 substances. Depending on the taxonomic group of interest, these data covered information on chronic toxicity for 20 – 31% of the compounds (Table 3).

**Table 3.**
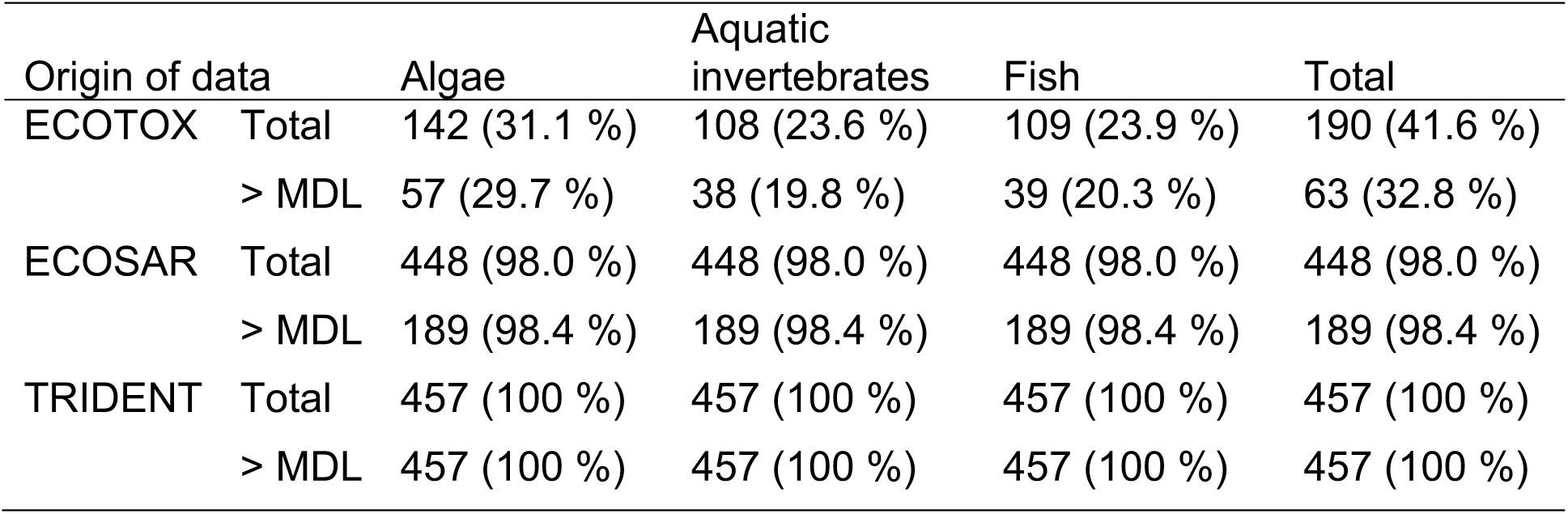
Overview on the chronic effect data per taxonomic group retrieved from the respective sources (ECOTOX database, ECOSAR and TRIDENT modelling) in the present study.

| Origin of data |  | Algae | Aquatic invertebrates | Fish | Total |
| --- | --- | --- | --- | --- | --- |
| ECOTOX | Total | 142 (31.1 %) | 108 (23.6 %) | 109 (23.9 %) | 190 (41.6 %) |
|  | > MDL | 57 (29.7 %) | 38 (19.8 %) | 39 (20.3 %) | 63 (32.8 %) |
| ECOSAR | Total | 448 (98.0 %) | 448 (98.0 %) | 448 (98.0 %) | 448 (98.0 %) |
|  | > MDL | 189 (98.4 %) | 189 (98.4 %) | 189 (98.4 %) | 189 (98.4 %) |
| TRIDENT | Total | 457 (100 %) | 457 (100 %) | 457 (100 %) | 457 (100 %) |
|  | > MDL | 457 (100 %) | 457 (100 %) | 457 (100 %) | 457 (100 %) |

ECOSAR was able to predict the chronic toxicity of 448 out of 457 compounds for all taxonomic groups of interest. For nine substances, the programme could not predict toxicity as these were found to be outside of the applicability domain (compounds with a log *K*_OW_ > 8 and/or a molecular weight > 1000 g/L are considered outside the applicability domain). As the TRIDENT model does have a larger applicability domain, we could retrieve chronic toxicity predictions for all taxonomic groups and all 457 compounds of interest. A representation of the relationships between the effect data sets including only the overlaps (i.e. showing for how many compounds we found empirical data from each taxonomic group as well as from *in silico* predictions) is depicted in an Euler diagram in Figure S1.

### 3.3. Mixture risk assessment and sites at risk

We applied 27 possible risk scenario combinations (i.e. nine hazard scenarios × three exposure scenarios) for each sample, each taxonomic group as well as the most sensitive taxonomic group (MST approach), resulting in a total of 756 risk quotient (RQ) sums (Figure S2).

To identify sites at risk, we considered a risk quotient sum in the MST approach (RQ_MST_) with a value equal to or higher than 1.0 as a threshold. Figure 3 represents a summary of the RQ_MST_ calculated for all samples and the five main hazard scenarios HS1 – HS5, which are detailed in Table 2. The *in silico* predictions for the remaining hazard scenarios HS2a, HS3a, HS2b and HS3b were multiplied and divided by a factor of 100 to examine model inaccuracies and to identify potential risk drivers. All risk quotients of the possible combinations of hazard and exposure scenarios are presented in Figure S2.

**Figure 3.**
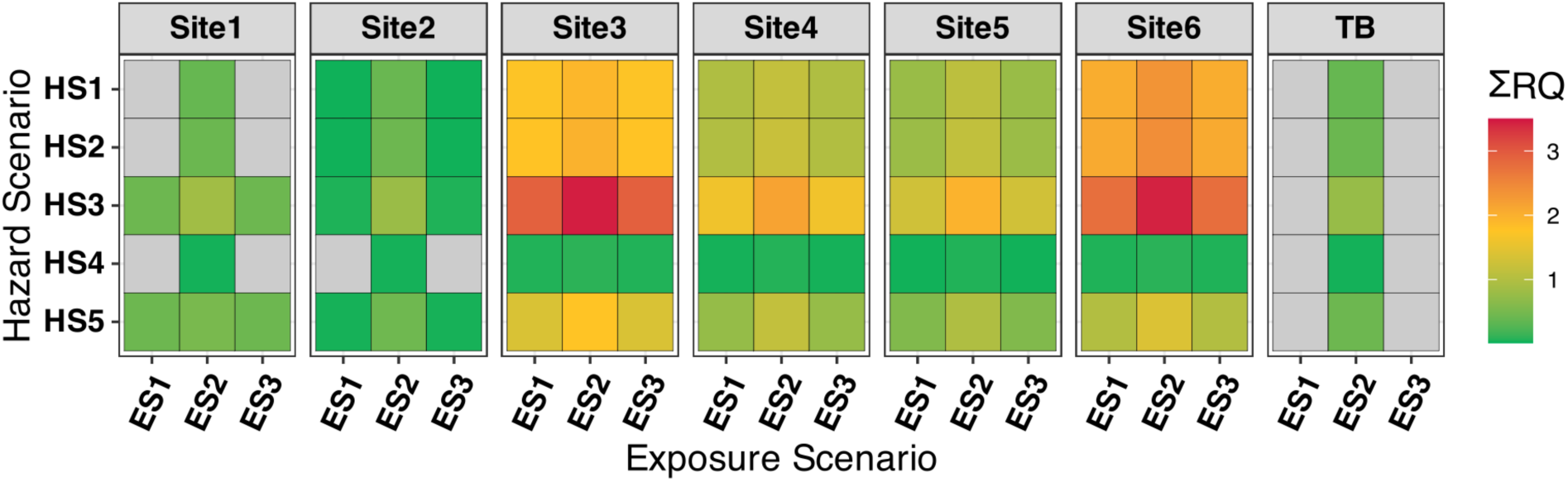
Summarising tile plot showing the Risk Quotient sums of the most sensitive taxonomic group (RQ_MST_) for each Exposure Scenario (ES) and the main five Hazard Scenarios (HS) and each sample (Site1-Site6). Note: the scale is continuous from 0.01 to 3.5 and grey tiles indicate an RQ sum below 0.01. TB: trip blank.

The lowest RQ_MST_ values were calculated for ES1 followed by ES3. Logically, using ES2 (all non-detects set to MDL) resulted in the highest RQ_MST_. A comparison of the individual MDLs with the obtained chronic toxicity data (Figure S3) indicates that the chemical-analytical methods used have a sufficiently high sensitivity in relation to the chronic toxicity of the compounds under consideration. Moreover, as previously observed by Inostroza et al. (2024), the inclusion of Kaplan-Meier estimates had only a small influence on the MRA outcome. This is also indicated by the minimal deviations between ES1 and ES3 (ΔRQ_MST_ ≤ 0.01 for HS1-HS3) in Figure S2. Consequently, we used the Kaplan-Meier estimation (ES3) as the most realistic exposure scenario to categorise sites as well as taxonomic groups at risk (Figure 4) and ES1 to identify mixture risk drivers.

**Figure 4.**
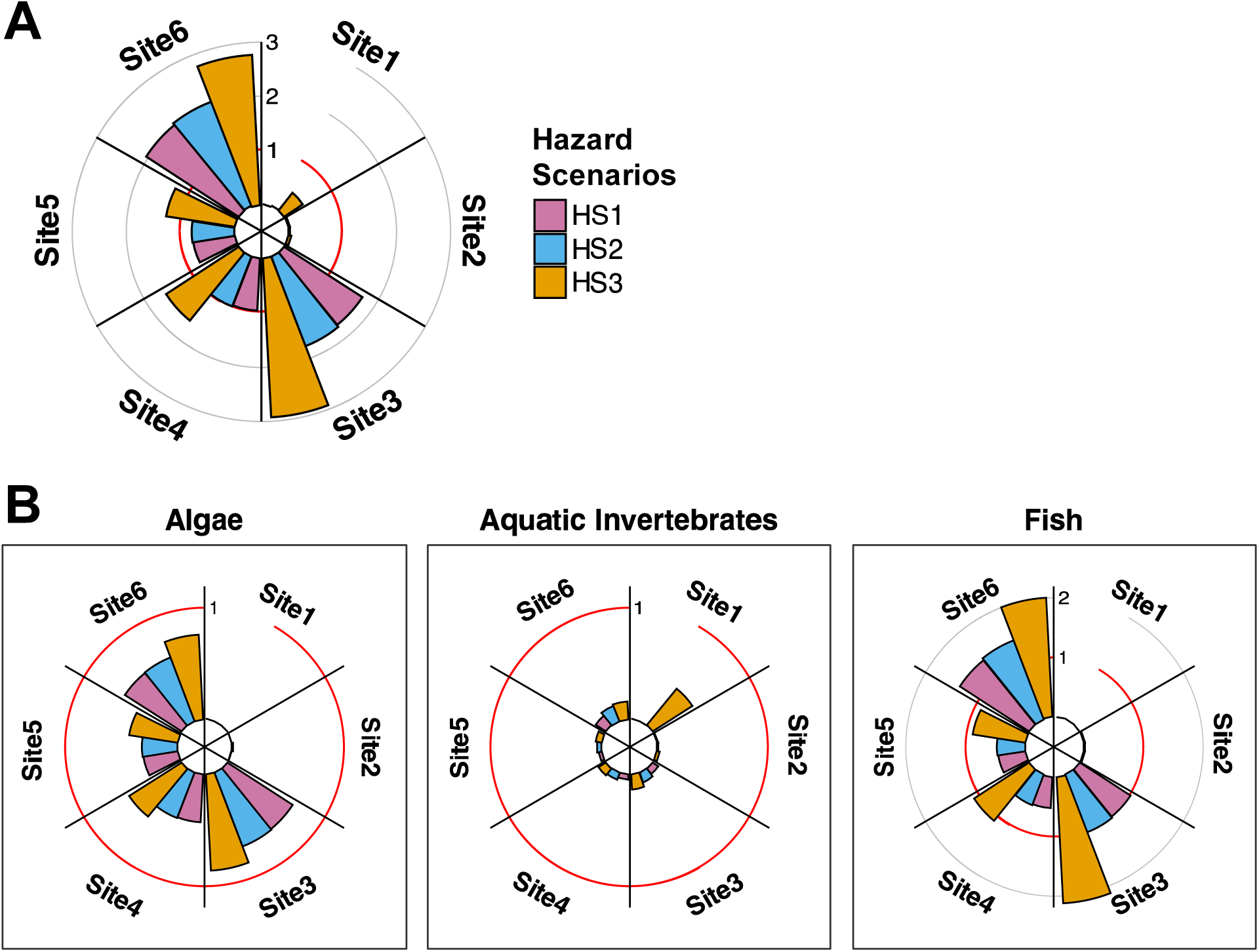
Circular bar plot showing the sum of Risk Quotients (RQs) for Exposure Scenario 3 and the three different Hazard Scenarios (HS) at each sampling site (Site1 – Site6). Panel (A) shows the results for the sum of RQs of the most sensitive trophic group. Panel (B) shows the results for the sum of RQs for each of the three taxonomic groups, algae, aquatic invertebrates and fish. Note that the y-axis in panel B is scaled differently to enhance readability.

HS1 resulted in the lowest, HS2 in slightly higher and HS3 in the highest RQ_MST_, as the RQs depend on the number of considered compounds when using the CA concept. For HS4 and HS5, the latter always resulted in higher risk estimates, though independent of the exposure scenario, the RQ_MST_ for the samples from Site3 – Site6 were lowest using HS4 and highest using HS3 (HS4<HS5<HS1<HS2<HS3). The results for the samples with fewer compounds detected (i.e. Site1, Site2 and the trip blank) indicated greater dependency between the hazard scenarios and the exposure scenarios on the resulting RQ_MST_.

Figure 4A shows that for the MST approach, Site3 and Site6 were identified as at risk independent of the underlying hazard scenario and using HS3 likewise indicated Site4 and Site5 as at risk. Nonetheless, when HS1 and HS2 were applied, the RQ_MST_ for Site4 (0.97 and 0.99, respectively) and Site5 (0.77 and 0.79, respectively) were close to the threshold value of 1.0. Figure 4B shows that none of the sites were indicated as at risk for the taxonomic group of algae, with the highest RQs at Site3 and Site6 ranging between 0.70 and 0.87 as well as 0.60 and 0.76, respectively. Similarly, the taxonomic group of aquatic invertebrates was not categorised at risk at any of the sampling sites with RQs between < 0.01 and 0.11 (HS1), < 0.01 and 0.14 (HS2) as well as 0.03 and 0.42 (HS3). Independent of the hazard scenario used, fish were indicated as a taxonomic group at risk at Site3 (RQs: 1.01 to 2.13) and Site6 (RQs: 1.38 to 2.02). Furthermore, the RQs for Site4 and Site5 exceeded the risk threshold or were close to it (1.09 and 0.89, respectively), when HS3 was applied.

### 3.4. Actual mixture risk drivers

To identify actual mixture risk drivers (i.e., compounds with an RQ_i_ ≥ 0.02 at any of the sites), we combined ES1 (non-detects set to zero) with HS3 (empirical toxicity data amended with *in silico* predictions from TRIDENT) and used the MST approach. The choice of HS3 as the most relevant hazard scenario for risk driver identification was made based on the fact that the TRIDENT model showed higher accuracy in predicting chronic toxicity for all taxonomic groups (Figure S4) compared to ECOSAR and is further elaborated in the discussion section.

We identified 21 compounds as actual mixture risk drivers (Figure 5, Table S3). These comprised 15 pharmaceuticals of various classes such as veterinary pharmaceuticals (ivermectin B1a), antibiotics (clarithromycin), anti-depressants/anxiolytics and antiepileptics (amitriptyline, citalopram, carbamazepine, temazepam, lamotrigine), diuretics (hydrochlorothiazide, furosemide), drugs targeting the cardio-vascular system (bezafibrate, metoprolol) as well as anaesthetics/analgesics and anti-inflammatory drugs (diclofenac, lidocaine, N-formyl-4-aminoantipyrine). The remaining six compounds are biocides (fipronil and hexadecylpyridinium) and pesticides (flumioxazin, propiconazole, azoxystrobin, dimethachlor CGA369873 and 2-hydroxydesethylterbuthylazine).

**Figure 5.**
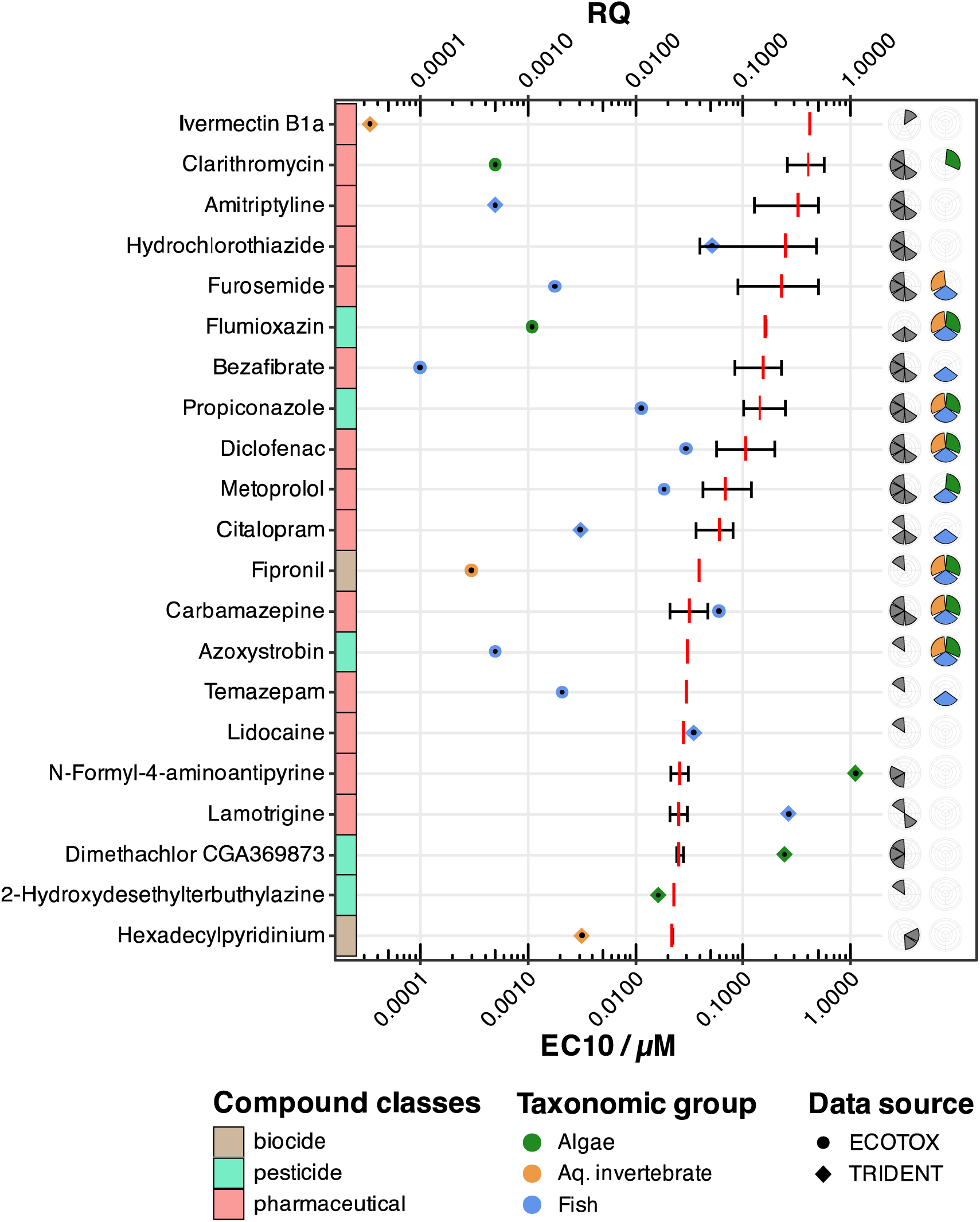
Illustration of the **actual** mixture risk drivers identified in the present study. The arithmetic mean risk quotient (RQ) of the compound is marked by the red vertical dash, and the whiskers indicate the minimum and maximum RQ. The colour of the tile adjacent to the compound name shows the compound class. The dots (empirical data from ECOTOX database) and diamonds (in silico prediction from TRIDENT), coloured depending on the taxonomic group, represent the most sensitive EC_10_-value used to calculate the RQs. The first column of pie charts specifies at which sampling site, the compound is identified as a risk driver (Site1, Site2, Site3, Site4, Site5, and Site6 on 1, 3, 5, 7, 9, and 11 o’clock, respectively). The second column of pie charts represents the presence and absence of empirical data for the three taxonomic groups, according to the same colour scheme as used for the EC_10_-values. Note: the x-axis is log-scaled and is representative of both the RQs and the EC_10_-values in µM.

For 12 of the actual risk drivers, we obtained empirical chronic toxicity data from the ECOTOX database for at least one of the taxonomic groups and for six, we were able to acquire empirical data for all taxonomic groups. Eleven risk drivers were identified based on empirical data, as the most sensitive EC_10_-equivalent was retrieved from the ECOTOX database. Most of the identified compounds were, in particular, risk drivers for the taxonomic group of fish, followed by algae and aquatic invertebrates. The EC_10_-equivalents for fish ranged from 0.9•10^-4^ µM (bezafibrate, ECOTOX) to 0.27 µM (lamotrigine, TRIDENT), for algae from 5.3•10^-4^ µM (clarithromycin, ECOTOX) to 1.13 µM (N-formyl-4-aminoantipyrine, TRIDENT), and for aquatic invertebrates from 0.3•10^-4^ µM (ivermectin B1a, TRIDENT) to 3.1•10^-3^ µM (hexadecylpyridinium, TRIDENT).

Of the total 21 actual risk drivers for the Holtemme River, 15 were identified at least at two sites, 11 in samples of at least three sites, 9 in samples of at least four sites, and five in samples from only one site.

### 3.5. Potential mixture risk drivers

For the identification of potential mixture risk drivers, we obtained data from the combination of the exposure scenario ES1 (non-detects set to zero) with hazard scenario HS3b (empirical toxicity data amended with *in silico* predictions from TRIDENT that were divided by a factor of 100) in the MST approach. With this analysis, we identified a total of 66 compounds flagged as potential mixture risk drivers. The full list of compounds is enclosed in Table S4 in the supplementary materials. Here, the focus will be set on the top 25 risk-driving compounds.

We identified 25 compounds from diverse compound classes (Figure 6), such as pesticides, dyes, polymer additives, personal care and household products, as well as pharmaceuticals, representing the largest share. These comprise an antiepileptic (lamotrigine), anaesthetics and analgesics (lidocaine, N-formyl-4-aminoantipyrine, tramadol, ketamine, 4-aminoantipyrine), antihypertensives (telmisartan, bisoprolol), an antipsychotic (pipamperone), an anti-androgen (bicalutamide), diuretics (torasemide, furosemide), and an antidiabetic (metformin). All of the most sensitive EC_10_-equivalents used for this risk scenario were provided by TRIDENT data. Most of the identified potential mixture risk drivers (11) were compounds exhibiting the lowest EC_10_ for fish, followed by algae (9) and aquatic invertebrates (5). The values for fish ranged from 3.5•10^-4^ µM (lidocaine) to 7.2•10^-2^ µM (5-methyl-1H-benzotriazole), for algae from 1.2•10^-4^ µM (trifloxystrobin NOA413161) to 0.14 µM (2-benzothiazolesulfonic acid) and for aquatic invertebrates from 2.6•10^-4^ µM (bicalutamide) to 3.6•10^-2^ µM (phenylbenzimidazole sulfonic acid).

**Figure 6.**
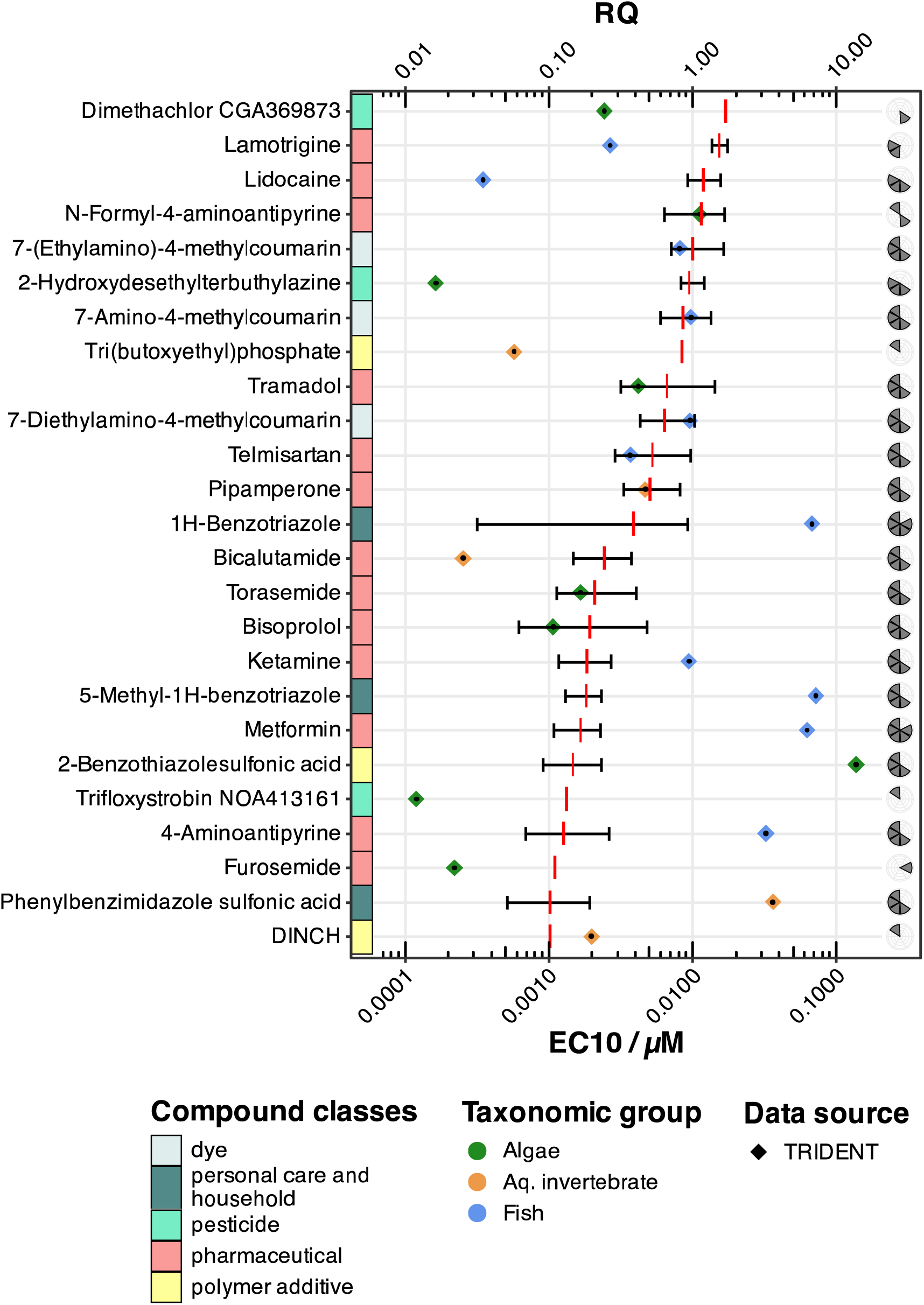
Illustration of the **potential** mixture risk drivers identified in the present study with a mean risk quotient (RQ) > 0.10. The mean RQ of the compound is marked by the red vertical dash, the whiskers indicate the minimum and maximum RQ. The colour of the tile adjacent to the compound name shows the compound class. The diamonds (in silico prediction from TRIDENT), coloured depending on the taxonomic group, represent the most sensitive EC_10_-value used to calculate the RQs. The column of pie charts specifies at which sampling site, the compound is identified as a risk driver (Site1, Site2, Site3, Site4, Site5, and Site6 on 1, 3, 5, 7, 9, and 11 o’clock, respectively). Note: the x-axis is log-scaled and is representative of both the RQs and the EC_10_-values in µM.

Six of the potential risk-driving compounds have already been identified as actual risk drivers in the present study (lamotrigine, lidocaine, N-formyl-4-aminoantipyrine, 2-hydroxydesethylterbuthylazine, dimethachlor CGA369873, furosemide). This is attributed to the fact that these compounds were identified as actual mixture risk drivers for at least one additional sites as compared to the actual mixture risk driver assessment. In total, 20 of the top 25 potential risk drivers for the Holtemme River were posing a risk at a minimum of two sites, 18 were identified in the samples from at least three sites, 16 at four sites and two compounds exceeded the threshold at five sites.

## 4. Discussion

We conducted a longitudinal chronic mixture risk assessment (MRA) using state-of-the-art large-volume water sampling techniques, LC-HRMS, and next-generation AI-based ecotoxicity modelling. To the best of our knowledge, this is the first use of deep learning architectures in chronic MRA combined with a real-world case study scenario. As discussed below, we identified hot spots of chemical contamination, highlighting priority sites for further investigation and/or risk mitigation. In addition, we identified fish as a particularly vulnerable taxonomic group and we highlighted the mixture risk drivers.

### 4.1. Exposure situation – complex mixtures of organic contaminants

The concentration and number of organic contaminants increased in the direction of the river flow and showed a clear pollution gradient, i.e. an increase in anthropogenic influence on the Holtemme River. Especially, the samples taken from downstream of the two WWTPs Silstedt and Halberstadt show that WWTPs are the main pathways for the introduction of complex chemical mixtures into the aquatic environment (Altenburger et al., 2019; Wolfram et al., 2021), each showing distinct pollution patterns (Figure 2A). The substances with the highest concentrations were compounds which are regularly found in surface waters (Finckh et al., 2024) but also compounds that are rather river basin specific such as the three compounds 7-diethylamino-4-methylcoumarin and its transformation products 7-ethylamino-4-methylcoumarin and 7-amino-4-methylcoumarin (Muschket et al., 2021, 2018; Weitere et al., 2021). In general, the concentration levels found are comparable to those of previous studies from the Holtemme River (Inostroza et al., 2017; Muschket et al., 2021; Švara et al., 2021; Tousova et al., 2017; Weitere et al., 2021) as well as to those of other European (Finckh et al., 2024, 2022; Loos et al., 2013) and non-European rivers (Carpenter and Helbling, 2018; Kandie et al., 2020) that are influenced by anthropogenic sources, specifically WWTPs.

### 4.2. Hazard quantification – chronic ecotoxicity data

Even though we have the technical capabilities and capacities to analyse numerous contaminants in environmental samples, we are still confronted with a lack of empirical data on the toxicity of those contaminants, especially when chronic toxicity is taken into account. This gap may be bridged by using QSAR approaches, which is also pointed out in guidance documents and reports on risk assessment of the European Chemicals Agency and the US EPA (European Chemicals Agency, 2020, 2008; US EPA, 1998), but there is concern in the scientific community that the scarcity of empirical data and the unreliability of QSAR prediction (e.g., domain of applicability, data scarcity and limited model validation) hinders a solid MRA (Gustavsson et al., 2023). Additionally, regulatory advisors recommend to not solely rely on QSARs for deriving environmental quality standards as they tend to underestimate toxicity (European Commission, 2018; Mark et al., 2008). Therefore, the generation of empirical data, as well as the development of new *in silico* methods, are indispensable (Gustavsson et al., 2024, 2023).

The ECOTOX database, one of the most comprehensive databases on ecotoxicological data, holds information on approx. 12,000 chemicals in more than 13,000 species, yielding over 1.1 Mio results (Olker et al., 2022). Nonetheless, data on chronic toxicity is still scarce as we were only able to obtain data on 32.8 % of the compounds quantified above the MDL for at least one of the taxonomic groups and only for 33 compounds (i.e. 17.2 %) for all taxonomic groups of interest (Table 3, Figure S1). To fill in the missing effect data, we derived values from *in silico* predictions (i.e. ECOSAR and TRIDENT) and were able to gather modelled ecotoxicological data for nearly all target substances. ECOSAR failed to predict toxicity data for eight compounds (amiodarone, DINCH, hexadecylpyridinium, montelukast, orlistat, salinomycin, telmisartan, tris(2-ethylhexl)phosphate) with predicted log *K*_OW_ values between 8.19 and 9.52, and one additional compound (vancomycin) with a molecular weight of 1449 g/mol since these compounds fell outside of the domain of applicability. When comparing the correlation between empirical and *in silico* data derived by ECOSAR and TRIDENT, it was demonstrated that the predictions by the TRIDENT model showed higher correlation coefficients for all taxonomic groups of interest (Figure S4). Interestingly, for fish, the correlation between ECOSAR and ECOTOX data was indicated as non-significant (α = 0.05) while TRIDENT to ECOTOX showed a highly significant correlation (Spearman’s correlation coefficient R = 0.67). Consequently, we concluded that the TRIDENT predictions more accurately line up with available empirical data and thus that TRIDENT predictions outperform ECOSAR. This is in agreement with the results of Gustavsson et al. (2024), specifically with respect to the prediction of chronic toxicity. Hence, we decided to use the hazard scenario HS3 (i.e. ECOTOX data amended by TRIDENT data) and HS3b (i.e. ECOTOX data amended by TRIDENT data divided by 100) as the most representative for the calculation of RQ sums (combined with ES3) and the identification of actual and potential mixture toxicity risk drivers (combined with ES1).

### 4.3. Mixture risk assessment

The systematic analysis and evaluation of 27 possible risk scenario combinations with three distinctive taxonomic groups from different trophic levels, as well as the MST approach, resulted in a detailed MRA for the Holtemme River. We identified four sites at risk (i.e., an RQ sum ≥ 1.0, depending on the risk scenario). Even with the use of the most conservative approach (non-detects set to zero and only empirical data used) the two sites downstream of the two WWTPs Silstedt and Halberstadt (Site3 and Site6) were flagged as being at risk, exceeding the RQ threshold by a factor of 1.73 and 2.04, respectively. This analysis assumes implicitly that neither other compounds (e.g., organic chemicals not included in the target list or metals) nor that those compounds lacking empirical data add to the toxicity of the sample. Even though CA might overestimate the risk of a given mixture slightly (Backhaus and Faust, 2012), this is likely not sufficient to account for the risk contribution of mixture components not accounted for during the assessment. This is particularly evident with regard to the empirical data scarcity of pharmaceuticals (Oelkers and Floeter, 2019; Spilsbury et al., 2024), as these active substances are designed to interact with human proteins and drug-target orthologues are evolutionarily conserved among various taxa (Gunnarsson et al., 2019). Consequently, and in connection with what was discussed above, we consider amending empirical data with *in silico* predictions a “more realistic” hazard scenario (HS3). This shows that all sites downstream of the first WWTP were exceeding the RQ threshold of 1 (Site3: 2.93, Site4: 1.61, Site5: 1.28, Site6: 2.77).

We applied a new approach in our assessment, as outlined in Inostroza et al. (2024), to determine various types of mixture risk drivers. The analysis showed that most of the actual and potential mixture risk drivers are pharmaceuticals. These pharmaceuticals are known to exhibit various target and non-target modes of action. While some are connected to antibiotic or anti-inflammatory properties, the largest set of the pharmaceuticals flagged as risk drivers are known neuroactive and cardio-vascular regulating substances (Kramer et al., 2024), such as amitriptyline, citalopram, temazepam, lamotrigine and hydrochlorothiazide as well as metoprolol. Furthermore, endocrine disrupting chemicals (EDCs) were found to be relevant mixture risk drivers, here specifically the known anti-androgenic dyes 7-ethylamino, 7-amino- and 7-diethylamino-4-methylcoumarin (Muschket et al., 2018) and bicalutamide (Kramer et al., 2024) were highlighted in our analysis. Interestingly, the anti-androgenic character was also quantified for the respective samples from Site3, Site4, Site5 and Site6, using a reporter gene assay (Weichert et al., 2024a).

Besides pharmaceuticals, pesticides are often described as important mixture risk drivers (Beckers et al., 2018; Malaj et al., 2014). The assessment in the present study showed that pesticides are the second-largest compound class among the risk drivers. In total, five parent compounds and eight transformation products were identified, which may be explained by the date of sampling. The timing of sampling during the end of October may introduce a degree of bias, as peak exposure events (such as the field application or spills of pesticides) cannot be accurately captured. However, the sampling method employed in the present study exemplifies an improvement in the ability to obtain a more chronic exposure representation, which is not feasible with grab sampling or the extraction of small volumes of surface water. Additionally, the seasonal timing of the study likely explains the predominant identification of transformation products rather than parent compounds of pesticides.

The seasonal aspect of the sampling is also likely to be responsible for why we identified fish as the most sensitive taxonomic groups. It has been shown that fish are more likely to be affected by chronic exposure, whereas algae and crustaceans are more likely to be affected by peak exposure and seasonal exposure (Beckers et al., 2018). This also follows the same pattern of evidence where we found pharmaceuticals to be the main drivers of toxicity, as known drug targets in humans have orthologues in fish (Gunnarsson et al., 2019).

## 5. Conclusion

We demonstrated an approach for an extensive AI-aided MRA, using a real-world case study scenario which opens new opportunities for future risk assessment strategies. Even though a component-based risk assessment, as presented in this study, is potentially drawn to overestimation of risk (Backhaus and Faust, 2012), our analyses using multiple exposure and hazard scenarios strongly suggest that chronic risk to aquatic organisms posed by exemplary environmental exposure scenarios tend to be rather underestimated with traditional risk assessment approaches. Moreover, we were able to identify pharmaceuticals as the most important risk-contributing chemical class, and fish to be the most sensitive taxonomic group, likely to be affected by chronic exposure during periods in which pesticide applications are low. Though we were able to obtain empirical ecotoxicological data on most of the identified mixture risk drivers for at least one of the taxonomic groups, the *in silico* predictions used in this study were particularly relevant for the evaluation of whole mixtures. This is of great importance to minimise the uncertainties connected to the lack of empirical (eco)toxicological data. Consequently, we strongly encourage the use of deep learning architectures, and multi-scenario approaches to consider uncertainties in risk assessment as well as the calculation of various types of risk drivers as established in the present study in future MRA.

Thus far the environmental impact following chronic exposure to multiple chemical stressors is largely unknown and the true risk is challenging to assess without the application of effect-oriented *in situ* studies (Altenburger et al., 2019; Backhaus et al., 2019). Therefore, we see great potential in future field experiments, combining a comprehensive MRA and involving e.g. fish to confirm or refute the genuine environmental impact of complex chemical mixtures.

## CRediT authorship contribution statement

**FG. Weichert**: Conceptualization, Data curation, Formal analysis, Investigation, Methodology, Software, Validation, Visualization, Writing – original draft, Writing – review & editing, Funding acquisition; **P. Inostroza**: Conceptualization, Data curation, Formal analysis, Investigation, Methodology, Software, Validation, Visualization, Writing – original draft, Writing – review & editing, Supervision; **J. Ahlheim**: Conceptualization, Methodology, Writing – review & editing; **T. Backhaus**: Conceptualization, Data curation, Formal analysis, Investigation, Methodology, Software, Validation, Writing – review & editing; **W. Brack**: Conceptualization, Writing – review & editing; **M. Brauns**: Conceptualization, Writing – review & editing; **P. Fink**: Conceptualization, Writing – review & editing; **M. Krauss**: Data curation, Formal analysis, Investigation, Writing – review & editing; **P. Svedberg**: Data curation, Formal analysis, Investigation, Methodology, Software, Validation, Writing – review & editing; **H. Hollert**: Conceptualization, Writing – review & editing, Supervision, Funding acquisition.

## Acknowledgments

We would like to extent our gratitude to Jakob Pfefferle and Margit Petre for invaluable support with the fieldwork, LVSPE set-up and extract preparation. The QExactive Plus LC-HRMS used at UFZ is part of the major infrastructure initiative CITEPro (Chemicals in the Terrestrial Environment Profiler) funded by the Helmholtz Association

## Supplementary Materials

**Figure S1.**
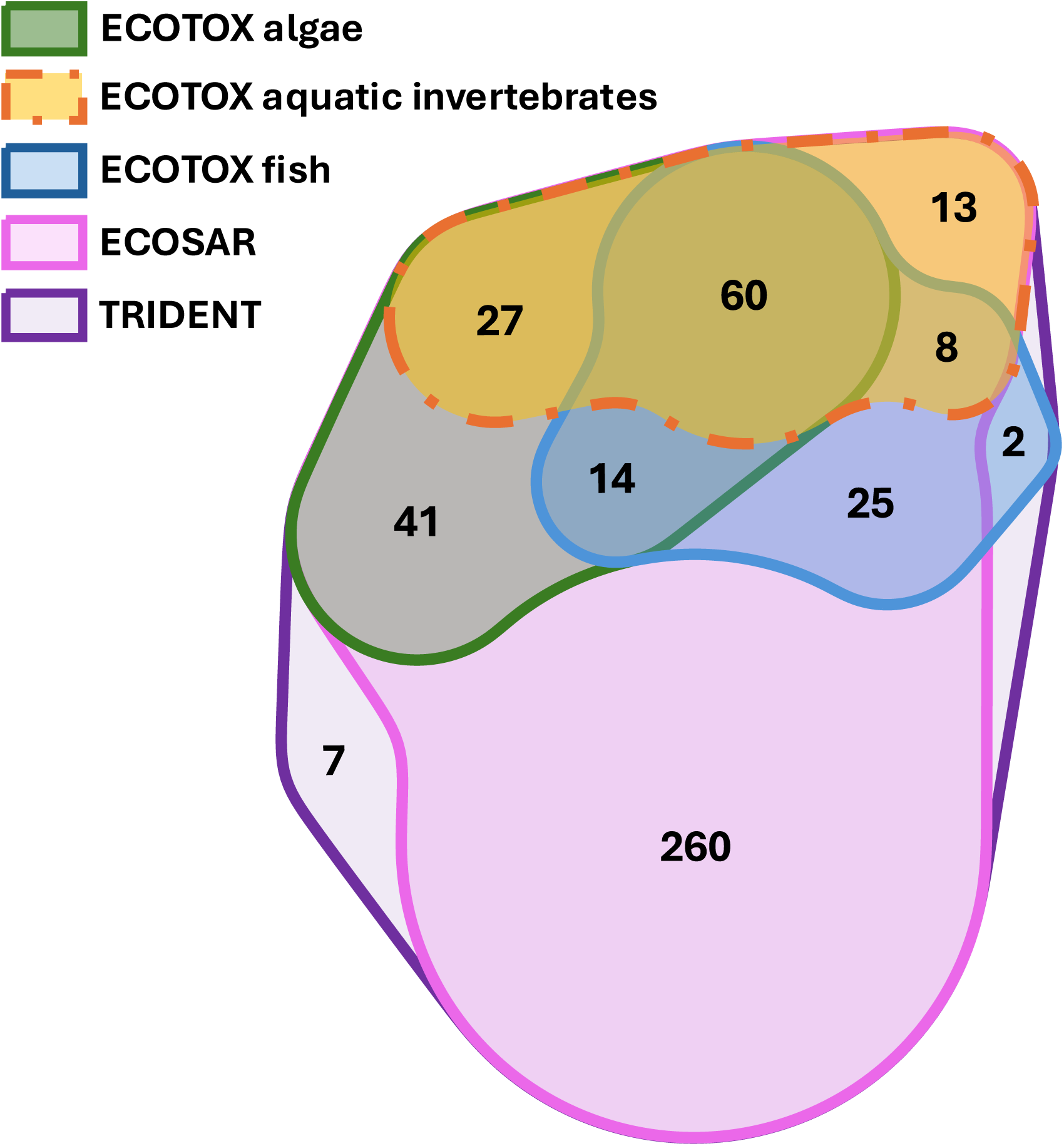
Quasi-proportional Euler diagram showing the relationship between the sets of effect data used in the present study. For example, the diagram shows that for 60 compounds we found empirical data from all taxonomic groups as well as from both in silico prediction tools. In addition, for 267 compounds we found in silico predictions only, of which 260 were predicted by both ECOSAR and TRIDENT and 7 were predicted by TRIDENT only. Note: The taxonomic groups were split only for empirical data, as the in silico models were able to predict toxicity independently of the taxonomic group of interest.

**Figure S2.**
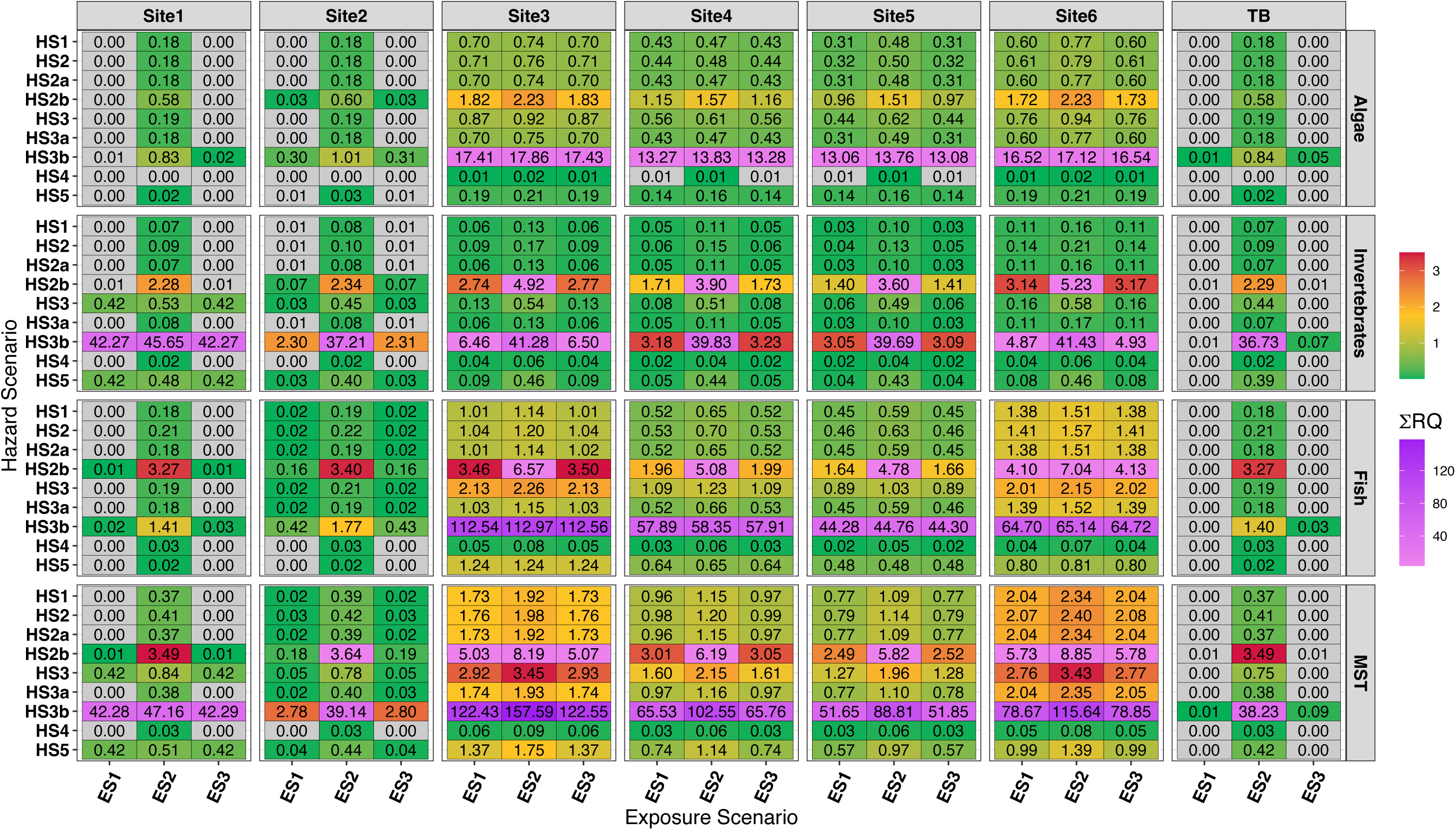
Tile plot showing the summed Risk Quotients (ΣRQ) for each sample (Site1 – Site6 and trip blank), calculated for algae, aquatic invertebrates, fish and the most sensitive trophic group for each Exposure Scenario (ES) and each Hazard Scenarios (HS). Note: the scale is continuous from 0.01 to 3.5 (green via yellow to red – the same colour code as in Figure 3) as well as from 3.51 to 158 (purple to violet) and grey tiles indicate a RQ sum below 0.01.

**Figure S3.**
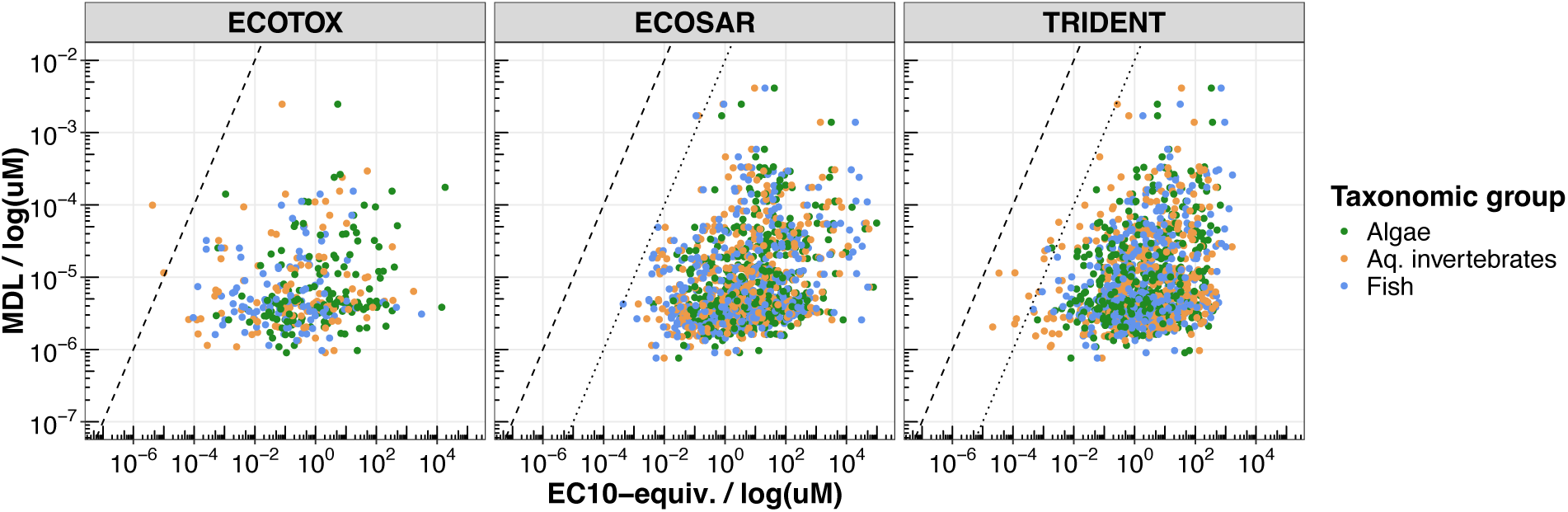
Scatter plot showing the relationship between method detection limit (MDL) and EC_10_-equivalents for each of the taxonomic groups from the three data sources ECOTOX, ECOSAR and TRIDENT. The dashed line represents the 1:1 line and the dotted line represents the theoretical 1:1 line when the in silico predictions are divided by a factor of 100 to account for modelling uncertainties.

**Figure S4.**
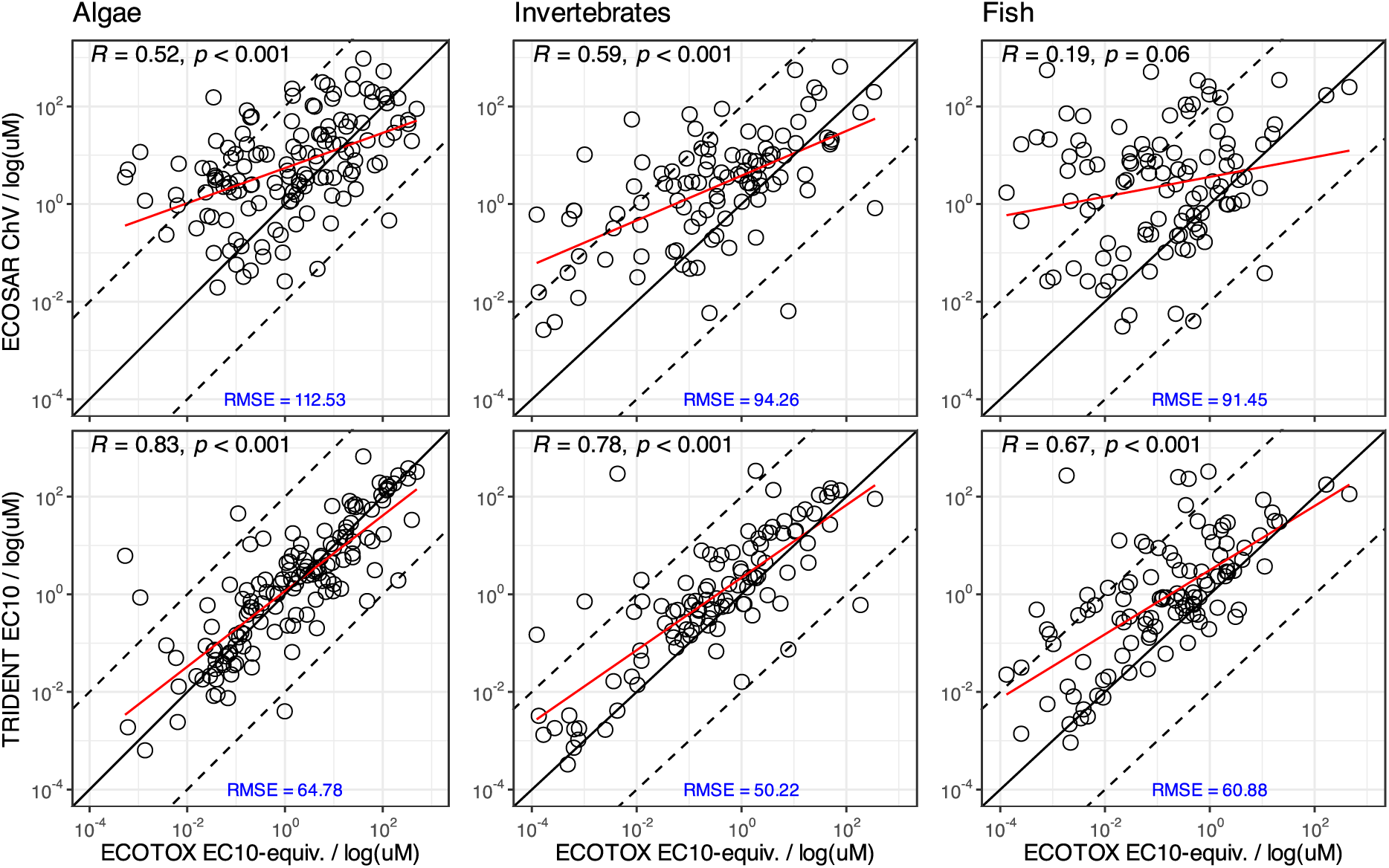
Spearman correlation analysis of empirical data vs. in silico predictions. The upper panel shows the correlation between empirical data and ECOSAR predictions (algae: n = 135, invertebrates: n = 104, fish: n = 104). The lower panel shows the correlation between empirical data and TRIDENT predictions (algae: n = 140, invertebrates: n = 106, fish: n = 108). R: correlation coefficient according to Spearman, RMSE: root mean squared error.

**Table S1.** Information on sampling sites (including coordinates, date and time of sampling as well as known effluent locations) and water parameters measured during the sampling. T: Temperature, EC: electrical conductivity.

| **Sampling site** | **km to river mouth** | **Long name** | **Coordinates (Long. / Lat.)** | **Date** | **Time** | **pH** | **T / °C** | **O<sub>2</sub> / mg/L** | **O<sub>2</sub> / %** | **EC / µS/cm** |
| --- | --- | --- | --- | --- | --- | --- | --- | --- | --- | --- |
| Site1 | 41.2 | MOBICOS Wernigerode / Steinerne Renne | 51.816980 / 10.724043 | 28.10.21 | 16:20 | 7.91 | 8.60 | 11.03 | 97.60 | 121.50 |
| Site2 | 34.1 | Urban Observatory 1 / upstream of WWTP Silstedt | 51.849829 / 10.797051 | 29.10.21 | 15:45 | 7.40 | 10.20 | 10.51 | 96.20 | 378.00 |
| - | 28.4 | Effluent WWTP Silstedt | 51.866342 / 10.863157 |  |  |  |  |  |  |  |
| Site3 | 27.7 | Urban Observatory 2 / downstream of WWTP Silstedt | 51.867888 / 10.873541 | 29.10.21 | 14:55 | 7.21 | 12.80 | 9.70 | 93.80 | 685.00 |
| Site4 | 19.5 | Gauging station Mahndorf | 51.885029 / 10.963073 | 30.10.21 | 10:11 | 7.25 | 9.60 | 10.67 | 95.30 | 706.00 |
| - | 11.8 | Effluent combined sewer overflow "Alter Sandfang" | 51.903052 / 11.058697 |  |  |  |  |  |  |  |
| Site5 | 11.5 | Upstream of WWTP Halberstadt downstream of combined sewer overflow | 51.904336 / 11.061795 | 29.10.21 | 10:47 | 7.05 | 9.50 | 11.11 | 98.90 | 745.00 |
| - | 11.5 | Effluent rain water retention basin "Alte Vorklärung" | 51.904395 / 11.061940 |  |  |  |  |  |  |  |
| - | 11.4 | Effluent WWTP Halberstadt | 51.904689 / 11.063058 |  |  |  |  |  |  |  |
| Site6 | 11.1 | Downstream of WWTP Halberstadt | 51.905442 / 11.067436 | 29.10.21 | 11:44 | 6.87 | 11.00 | 10.20 | 94.60 | 884.00 |

**Table S2.** The number of detected compounds above and below the respective method detection limit (MDL), as well as the number of non-detects, for each sample. *Compounds quantified below the MDL must be considered statistically not different from zero. TB: trip blank.

| Sample | > MDL | < MDL* | Non-detects |
| --- | --- | --- | --- |
| TB | 5 | 7 | 445 |
| Site1 | 8 | 6 | 443 |
| Site2 | 40 | 24 | 393 |
| Site3 | 157 | 32 | 268 |
| Site4 | 138 | 31 | 288 |
| Site5 | 126 | 35 | 296 |
| Site6 | 169 | 27 | 261 |
| Total | 192 | 42 | 223 |

**Table S3.** Actual mixture risk drivers as, identified from the MST approach (most sensitive taxonomic group) using the exposure scenario ES1 combined with the hazard scenario HS3. The table provides a summary of the compounds and their respective class, as well as the minimum (min), maximum (max), mean risk quotient, and number of sites at which the compound was identified as a risk-driving substance. Additionally, EC10-equivalents for empirical data (ECOTOX) and in silico predictions (TRIDENT) are presented for all taxonomic groups of interest, including algae, aquatic invertebrates (inv.) and fish.

| Compound | Compound class | Risk Quotient | | | # of sites | ECOTOX / $\mu\text{M}$ | | | TRIDENT / $\mu\text{M}$ | | |
| --- | --- | --- | --- | --- | --- | --- | --- | --- | --- | --- | --- |
|  |  | Min | Mean | Max |  | Algae | Inv. | Fish | Algae | Inv. | Fish |
| Ivermectin B1a | Pharmaceutical | 0.42 | 0.42 | 0.42 | 1 | - | - | - | 0.32 | $3 \cdot 10^{-5}$ | 0.10 |
| Clarithromycin | Pharmaceutical | 0.27 | 0.41 | 0.57 | 4 | $5 \cdot 10^{-4}$ | - | - | 6.15 | 3.30 | 0.32 |
| Amitriptyline | Pharmaceutical | 0.13 | 0.33 | 0.51 | 4 | - | - | - | 1.93 | 0.78 | $5 \cdot 10^{-4}$ |
| Hydrochlorothiazide | Pharmaceutical | 0.04 | 0.25 | 0.49 | 4 | - | - | - | 0.30 | 4.11 | 0.05 |
| Furosemide | Pharmaceutical | 0.09 | 0.23 | 0.51 | 4 | - | 30.23 | $2 \cdot 10^{-3}$ | 0.02 | 108.63 | 271.40 |
| Flumioxazin | Pesticide | 0.16 | 0.16 | 0.16 | 2 | $1 \cdot 10^{-3}$ | 0.10 | 1.41 | 0.86 | 0.71 | 2.34 |
| Bezafibrate | Pharmaceutical | 0.08 | 0.16 | 0.23 | 4 | - | - | $1 \cdot 10^{-4}$ | 14.94 | 0.22 | 323.47 |
| Propiconazole | Pharmaceutical | 0.10 | 0.15 | 0.25 | 4 | 0.70 | 0.49 | 0.01 | 1.02 | 0.75 | 1.36 |
| Diclofenac | Pharmaceutical | 0.06 | 0.11 | 0.20 | 4 | 7.66 | 0.67 | 0.03 | 1.94 | 7.21 | 1.77 |
| Metoprolol | Pharmaceutical | 0.04 | 0.07 | 0.12 | 4 | 5.46 | - | 0.02 | 6.13 | 3.50 | 12.67 |
| Citalopram | Pharmaceutical | 0.04 | 0.06 | 0.08 | 3 | - | - | $5 \cdot 10^{-3}$ | 3.96 | 12.75 | $3 \cdot 10^{-3}$ |
| Fipronil | Biocide | 0.04 | 0.04 | 0.04 | 1 | 0.20 | $3 \cdot 10^{-4}$ | 0.03 | 0.54 | $2 \cdot 10^{-3}$ | 0.02 |
| Carbamazepine | Pharmaceutical | 0.02 | 0.03 | 0.05 | 4 | 1.23 | 0.09 | 0.06 | 3.11 | 0.19 | 9.78 |
| Azoxystrobin | Pesticide | 0.03 | 0.03 | 0.03 | 1 | 0.17 | 0.01 | $5 \cdot 10^{-4}$ | 1.22 | 0.02 | 0.49 |
| Temazepam | Pharmaceutical | 0.03 | 0.03 | 0.03 | 1 | - | - | $2 \cdot 10^{-3}$ | 10.05 | 24.31 | $2 \cdot 10^{-3}$ |
| Lidocaine | Pharmaceutical | 0.03 | 0.03 | 0.03 | 1 | - | - | - | 216.19 | 28.86 | 0.04 |
| N-Formyl-4-aminoantipyrine | Pharmaceutical | 0.02 | 0.03 | 0.03 | 2 | - | - | - | 1.13 | 3.28 | 3.78 |
| Lamotrigine | Pharmaceutical | 0.02 | 0.03 | 0.03 | 2 | - | - | - | 1.89 | 3.17 | 0.27 |
| Dimethachlor CGA369873 | Pesticide | 0.02 | 0.03 | 0.03 | 3 | - | - | - | 0.25 | 124.41 | 117.67 |
| 2-Hydroxy-desethylterbuthylazine | Pesticide | 0.02 | 0.02 | 0.02 | 1 | - | - | - | 0.02 | 5.47 | 0.14 |
| Hexadecylpyridinium | Biocide | 0.02 | 0.02 | 0.02 | 2 | - | - | - | 0.07 | $3 \cdot 10^{-3}$ | 0.19 |

**Table S4.** Potential mixture risk drivers as, identified from the MST approach (most sensitive taxonomic group) using the exposure scenario ES1 combined with the hazard scenario HS3b. The table provides a summary of the compounds and their respective class, as well as the minimum (min), maximum (max), mean risk quotient, and number of sites at which the compound was identified as a risk-driving substance. Additionally, EC10-equivalents for empirical data (ECOTOX) and in silico predictions (TRIDENT) are presented for all taxonomic groups of interest, including algae, aquatic invertebrates (inv.) and fish. It should be noted that the EC10-equivalents provided in the in silico predictions do not include the division by 100, as detailed in the materials and methods section.

| Compound | Compound class | Risk Quotient | | | # of sites | ECOTOX / μM | | | TRIDENT / μM | | |
| --- | --- | --- | --- | --- | --- | --- | --- | --- | --- | --- | --- |
|  |  | Min | Mean | Max |  | Algae | Inv. | Fish | Algae | Inv. | Fish |
| Dimethachlor CGA369873 | Pesticide | 1.71 | 1.71 | 1.71 | 1 | - | - | - | 0.25 | 124.41 | 117.67 |
| Lamotrigine | Pharmaceutical | 1.36 | 1.55 | 1.74 | 2 | - | - | - | 1.89 | 3.17 | 0.27 |
| Lidocaine | Pharmaceutical | 0.92 | 1.20 | 1.56 | 3 | - | - | - | 216.19 | 28.86 | 0.04 |
| N-Formyl-4-aminoantipyrine | Pharmaceutical | 0.64 | 1.16 | 1.68 | 2 | - | - | - | 1.13 | 3.28 | 3.78 |
| 7-(Ethylamino)-4-methylcoumarin | Dye | 0.71 | 1.01 | 1.65 | 4 | - | - | - | 15.57 | 6.34 | 0.82 |
| 2-Hydroxydesethylterbutylazine | Pesticide | 0.84 | 0.96 | 1.20 | 3 | - | - | - | 0.02 | 5.47 | 0.14 |
| 7-Amino-4-methylcoumarin | Dye | 0.60 | 0.87 | 1.35 | 4 | - | - | - | 29.50 | 16.42 | 0.98 |
| Tri(butoxyethyl)phosphate | Polymer additive | 0.86 | 0.86 | 0.86 | 1 | - | - | - | 12.76 | 0.06 | 2.65 |
| Tramadol | Pharmaceutical | 0.32 | 0.67 | 1.44 | 4 | - | - | - | 0.42 | 58.86 | 0.42 |
| 7-Diethylamino-4-methylcoumarin | Dye | 0.43 | 0.64 | 1.03 | 4 | - | - | - | 7.50 | 6.13 | 0.97 |
| Telmisartan | Pharmaceutical | 0.29 | 0.53 | 0.96 | 4 | - | - | - | 11.21 | 1.43 | 0.37 |
| Pipamperone | Pharmaceutical | 0.33 | 0.51 | 0.82 | 4 | - | - | - | 1.57 | 0.47 | 1.04 |
| 1H-Benzotriazole | Personal care and household | 0.03 | 0.39 | 0.93 | 5 | - | - | - | 122.87 | 75.76 | 6.80 |
| Bicalutamide | Pharmaceutical | 0.15 | 0.25 | 0.37 | 4 | - | - | 0.09 | 0.48 | 0.03 | 0.22 |
| Torasemide | Pharmaceutical | 0.11 | 0.21 | 0.41 | 4 | - | - | - | 0.17 | 156.72 | 23.92 |
| Bisoprolol | Pharmaceutical | 0.06 | 0.20 | 0.48 | 4 | - | - | - | 0.11 | 1.85 | 1.30 |
| Ketamine | Pharmaceutical | 0.12 | 0.19 | 0.27 | 4 | - | - | - | 9.72 | 2.80 | 0.95 |
| 5-Methyl-1H-benzotriazole | Personal care and household | 0.13 | 0.18 | 0.23 | 4 | - | - | - | 85.29 | 67.38 | 7.24 |
| Metformin | Pharmaceutical | 0.11 | 0.17 | 0.23 | 5 | - | - | - | 135.65 | 159.05 | 6.31 |
| 2-Benzothiazolesulfonic acid | Polymer additive | 0.09 | 0.15 | 0.23 | 4 | - | - | - | 13.81 | 26.13 | 925.54 |
| Trifloxystrobin NOA413161 | Pesticide | 0.13 | 0.13 | 0.13 | 1 | - | - | - | 0.01 | 2.78 | 0.08 |
| 4-Aminoantipyrine | Pharmaceutical | 0.07 | 0.13 | 0.26 | 4 | - | - | - | 25.18 | 11.09 | 3.26 |
| Furosemide | Pharmaceutical | 0.11 | 0.11 | 0.11 | 1 | - | 30.23 | 2•10<sup>-3</sup> | 0.02 | 108.63 | 271.40 |
| Phenylbenzimidazole sulfonic acid | Personal care and household | 0.05 | 0.10 | 0.19 | 4 | - | - | - | 92.03 | 3.64 | 91.20 |
| DINCH | Polymer additive | 0.10 | 0.10 | 0.10 | 1 | - | - | - | 38.07 | 0.20 | 9.27 |
| 10,11-Dihydro-10,11-dihydroxycarbamazepine | Pharmaceutical | 0.05 | 0.10 | 0.16 | 4 | - | - | - | 3.25 | 368.89 | 13.09 |
| Sucralose | Food and beverage | 0.07 | 0.09 | 0.12 | 4 | - | - | - | 49.74 | 23.43 | 54.29 |
| Gabapentin-Lactam | Pharmaceutical | 0.05 | 0.08 | 0.11 | 4 | - | - | - | 263.82 | 12.90 | 2.39 |
| Chlorophene | Biocide | 0.08 | 0.08 | 0.08 | 1 | - | - | - | 0.08 | 0.12 | 0.99 |
| Azithromycin | Pharmaceutical | 0.02 | 0.07 | 0.12 | 4 | - | - | - | 2.83 | 1.92 | 0.30 |
| Trifloxystrobin CGA 321113 | Pesticide | 0.04 | 0.06 | 0.08 | 4 | - | - | - | 0.01 | 0.09 | 0.06 |
| o-Dianisidine | Intermediate | 0.05 | 0.06 | 0.07 | 4 | 13.51 | - | - | 14.69 | 1.89 | 6.48 |
| TMDD | Additive | 0.04 | 0.06 | 0.09 | 4 | - | - | - | 3.91 | 7.77 | 64.73 |
| Tris(1-chloro-2-propyl)phosphate | Polymer additive | 0.04 | 0.06 | 0.10 | 5 | - | - | - | 99.28 | 1.32 | 5.75 |
| Valsartan | Pharmaceutical | 0.02 | 0.06 | 0.14 | 4 | - | - | - | 156.11 | 14.60 | 0.56 |
| 2-(Methylthio)benzothiazole | Polymer additive | 0.02 | 0.05 | 0.09 | 2 | - | - | - | 0.65 | 8.21 | 5.28 |
| Hexa(methoxymethyl)melamine | Polymer additive | 0.04 | 0.05 | 0.07 | 2 | - | - | - | 8.99 | 0.64 | 1.42 |
| Caffeine | Food and beverage | 0.05 | 0.05 | 0.05 | 1 | 494.36 | 0.42 | - | 322.76 | 0.55 | 3.15 |
| Triphenylphosphine oxide | Intermediate | 0.05 | 0.05 | 0.05 | 1 | - | - | - | 7.51 | 47.50 | 16.73 |
| Mirtazapine | Pharmaceutical | 0.02 | 0.05 | 0.08 | 4 | - | - | - | 0.78 | 3.41 | 0.72 |
| 2-Hydroxycarbamazepine | Pharmaceutical | 0.03 | 0.05 | 0.08 | 4 | - | - | - | 7.40 | 0.45 | 4.16 |
| 4-Hydroxybenzotriazole | Personal care and household | 0.04 | 0.05 | 0.05 | 2 | - | - | - | 149.82 | 70.68 | 5.47 |
| Ambroxol | Pharmaceutical | 0.05 | 0.05 | 0.05 | 1 | - | - | - | 1.88 | 0.50 | 1.40 |
| Azoxystrobin acid | Pesticide | 0.05 | 0.05 | 0.05 | 1 | - | - | - | 0.40 | 0.08 | 0.51 |
| Oxazepam | Pharmaceutical | 0.02 | 0.05 | 0.07 | 2 | - | - | - | 6.85 | 0.13 | 0.47 |
| Pindolol | Pharmaceutical | 0.03 | 0.05 | 0.06 | 5 | - | - | - | 1.06 | 1.28 | 0.27 |
| p-Toluenesulfonamide | Intermediate | 0.04 | 0.04 | 0.04 | 1 | - | - | - | 38.67 | 598.13 | 736.18 |
| Metoprolol acid | Pharmaceutical | 0.04 | 0.04 | 0.04 | 4 | - | - | - | 22.13 | 28.27 | 53.61 |
| Sotalol | Pharmaceutical | 0.02 | 0.04 | 0.06 | 2 | - | - | - | 0.30 | 47.29 | 93.36 |
| Diphenhydramine | Pharmaceutical | 0.04 | 0.04 | 0.04 | 1 | - | - | - | 13.74 | 4.13 | 16.70 |
| Dimethenamid ESA | Pesticide | 0.03 | 0.03 | 0.04 | 2 | - | - | - | 0.02 | 17.73 | 321.28 |
| Kresoxim-methyl | Pesticide | 0.03 | 0.03 | 0.03 | 1 | 0.10 | - | - | 0.15 | 0.13 | 0.04 |
| Fluconazole | Pharmaceutical | 0.03 | 0.03 | 0.04 | 4 | 10.00 | - | - | 8.92 | 2.86 | 0.53 |
| Candesartan | Pharmaceutical | 0.03 | 0.03 | 0.04 | 2 | - | - | - | 8.62 | 153.89 | 44.16 |
| Carbetamide | Pesticide | 0.03 | 0.03 | 0.03 | 1 | - | - | - | 0.10 | 5.51 | 253.08 |
| 1,3-Diphenylguanidine | Polymer additive | 0.03 | 0.03 | 0.03 | 1 | - | - | - | 10.59 | 2.11 | 14.88 |
| Fluvoxamine | Pharmaceutical | 0.02 | 0.03 | 0.04 | 2 | 9.99 | - | - | 1.62 | 0.51 | 0.71 |
| Metolachlor-NOA 413173 | Pesticide | 0.03 | 0.03 | 0.03 | 2 | - | - | - | 8.09 | 1.58 | 177.31 |
| Prosulfocarb | Pesticide | 0.02 | 0.03 | 0.03 | 2 | - | - | - | 3.43 | 1.14 | 0.07 |
| Terbutryn | Biocide | 0.02 | 0.02 | 0.02 | 1 | 0.02 | - | 4.14 | 0.09 | 0.18 | 5.09 |
| Lorazepam | Pharmaceutical | 0.02 | 0.02 | 0.02 | 1 | - | - | - | 5.16 | 0.17 | 0.75 |
| 10,11-Dihydro-10-hydroxycarbamazepine | Pharmaceutical | 0.02 | 0.02 | 0.02 | 1 | - | - | - | 4.50 | 10.72 | 9.20 |
| Clozapine | Pharmaceutical | 0.02 | 0.02 | 0.02 | 1 | - | - | 0.07 | 0.77 | 0.40 | 0.13 |
| Phenazone | Pharmaceutical | 0.02 | 0.02 | 0.02 | 1 | - | - | - | 132.56 | 2.67 | 3.34 |
| Melperon | Pharmaceutical | 0.02 | 0.02 | 0.02 | 1 | - | - | - | 5.88 | 3.53 | 0.79 |
| Lauramidopropylbetaine | Personal care and household | 0.02 | 0.02 | 0.02 | 1 | - | - | - | 13.23 | 4.59 | 224.34 |

## Footnotes

1 EC: effect concentration, LC: lethal concentration, IC: inhibitory concentration, ED: effective dose, NOEC/NOEL: no-observed-effect concentration/level, LOEC/LOEL: lowest-observed-effect concentration/level, MATC: maximum acceptable toxicant concentration.

